# Household level insecticide deployments (micro-mosaics) for insecticide resistance management: Evaluating deliberate and accidental deployments

**DOI:** 10.1101/2024.07.05.602220

**Authors:** Neil Philip Hobbs, Ian Michael Hastings

**Author notes:** Corresponding Author &.

## Abstract

Mixtures of two insecticides in a single formulation at full-dose are frequently evaluated as the “best” insecticide resistance (IRM) strategy in public health. However, this requires both insecticides to be mixed together in a single formulation which may not be possible or practical. Deploying different insecticides in different households (“micro-mosaics”) may allow for mosquitoes to encounter different insecticides in subsequent gonotrophic cycles obtaining a “temporal mixture”. We evaluate micro-mosaics considering their deliberate use and their accidental use using a mathematical model assuming polygenic resistance. Deliberate micro-mosaics are evaluated against rotations and mixtures (full-dose or half-dose) over a range of scenarios allowing for cross resistance and insecticide decay on their ability to slow the development of resistance. Accidental micro-mosaics are evaluated to understand the implication of mixture insecticide-treated nets (ITNs) and standard (pyrethroid only) ITNs being deployed alongside one another on the development of resistance across a range of initial resistance scenarios. Deliberate micro-mosaics are found to not differ substantially in their IRM capability from either rotations or half-dose mixtures. When micro-mosaics do outperform rotations or half-dose mixtures the benefit is often small. Micro-mosaics are found to perform worse than full-dose mixtures. Accidental micro-mosaics are found to reduce the ability of mixtures to slow the development of resistance. The deployment of deliberate micro-mosaics was found to not be beneficial versus rotations or mixtures indicating this strategy should not be pursued. Micro-mosaics occurring accidentally due to multiple distribution channels inhibits the effectiveness of mixture ITNs in slowing the development of resistance. Where mixture ITNs are used keep the coverage of the mixture high relative to standard (pyrethroid-only) ITNs is key.

## 1. Introduction

Insecticide resistance (IR) is an issue for the control of vector-borne diseases (Hemingway et al., 2016). Mitigating the impact of IR may be possible by adopting insecticide resistance management (IRM) strategies which can be designed around the deployments of insecticides over space, time and dose (Rex Consortium, 2013). Often, emphasis is placed on evaluating the temporal deployment of insecticides such as comparing sequences, rotations, or mixture deployments. Mixtures have been found to be a highly promising IRM strategy (Curtis, 1985; Helps et al., 2017; Hobbs et al., 2023; Hobbs & Hastings, 2024; Madgwick & Kanitz, 2022; Mani, 1985; South & Hastings, 2018), providing the condition of high effectiveness is maintained (Rex Consortium, 2013). Unfortunately, this would require both insecticides in the mixture to be at their monotherapy application dose, inevitably increasing the cost of the mixture product.

The deployment of insecticides over space (mosaics), is less studied (Rex Consortium, 2010), although some more recent studies are including landscape simulations (Baudrot et al., 2023; Duthie et al., 2022; Hardy, 2022; Slater et al., 2017) albeit these simulation platforms are more applicable to agriculture than public health. One challenge in evaluating mosaics comes from the terminology used to describe mosaics, whereby the term the term “mosaic” is used regardless of the spatial scale of its deployments. In public health, the term mosaic has been used to describe insecticide deployments at the village or district level (WHO, 2012), down to the individual bed-net level where different net panels have different insecticides (Corbel et al., 2010). This again highlights an issue with evaluating IRM strategies and comparing between studies, in that deciphering each strategy is often unclear.

In public health, insecticides in the form of insecticide treated nets (ITNs) or indoor residual spraying (IRS) are distributed at the household level, but deployment decisions are made at larger spatial scales. That is, it could be technically possible for different insecticides to be given to different households in the same village during a distribution. This household level distribution of different insecticides has been termed micro-mosaics and the more landscape level distributions as macro-mosaics (Jones et al., 2023; Levick et al., 2017). A micro-mosaic is defined as the deployment of insecticides at a fine spatial scale such that the insecticides are present in the same intervention site (e.g., village) but are sufficiently spatially separated such that a mosquito would encounter only one insecticide at a time (Jones et al., 2023). For example, a female mosquito would encounter only one insecticide in each gonotrophic cycle. This is a distinction from an ITN and IRS combination or panel nets where it is possible for two insecticides to be present in the same household and therefore both insecticides may be encountered near simultaneously despite being spatially separated.

The proposed theoretical benefit of micro-mosaics is to obtain a “temporal mixture” from mosquitoes encountering different insecticides throughout their lifetime (Jones et al., 2023). It is hypothesised that this “temporal mixture” may allow some, or even all, the beneficial effects of a true mixture when the latter is itself unavailable due, for example, production by different manufacturers or the two insecticides not forming a chemically stable mixture formulation. Micro-mosaics would initially be hypothesised to perform between the effectiveness of rotations and full-dose mixtures. However, whether micro-mosaics perform more similarly to rotations or full-dose mixtures is unclear. This is especially important when considering cross resistance, where positive cross-resistance has been found to be especially detrimental for rotations and less detrimental for full-dose mixtures (Hobbs et al., 2023).

Micro-mosaics may be considered as both “deliberate” and “accidental” strategies (Figure 1). Deliberate micro-mosaics are the intentional use of micro-mosaics as part of a planned IRM strategy as implemented by vector control programmes. Deliberate micro-mosaics are expected to be a complex IRM strategy to implement, due to the requirement of multiple procurement and logistical pathways. Computer simulations are therefore required to identify, and quantify, the likely benefits of the deliberate micro-mosaic strategy before attempting costly trials of a new IRM strategy (Tabashnik, 1986).

**Figure 1:**
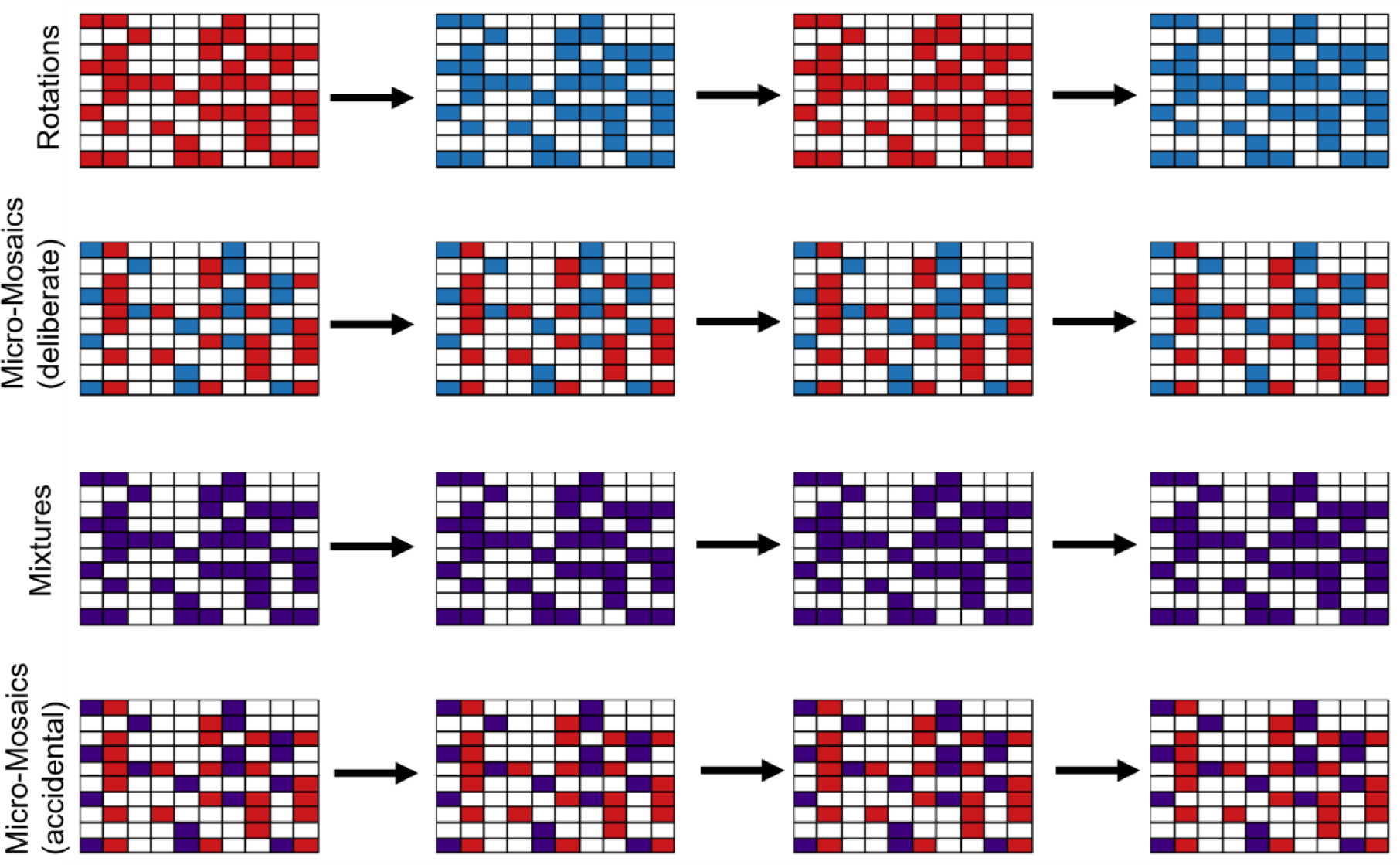
Diagrammatic representation of IRM strategies. Each tile represents either a house treated with insecticide (coloured) or a house untreated and or part of the village where the insecticide is not deployed (white). Rows represent IRM strategies and columns are sequential timesteps (e.g., consecutive transmission seasons). Top row is the rotation strategy, where the insecticides (*i* = blue, *j* = red) are deployed separately over time with the insecticides deployed rotated at each deployment interval. Second row is the deliberate micro-mosaics where a proportion of the households receive insecticide *i* (blue) and the other proportion receive insecticide *j* (red), mosquitoes may therefore encounter different insecticides in each gonotrophic cycle. The third row is the mixture strategy, where insecticides *i* (blue) and *j* (red) are in a single formulation of *ij* = purple) such that any mosquito exposed to the mixture is inevitably exposed to both insecticides. The bottom row is accidental micro-mosaics strategy whereby mixtures (*ij*= purple) and monotherapy applications (*j* = red) are deployed at the same time, such that some households have both *ij* (purple) and some have only *j* (red) as may occur with the deployment of next-generation mixture ITNs and standard pyrethroid ITNs.

Accidental micro-mosaics occur unintentionally because of insecticidal products having different distribution channels. For example, ITNs can be distributed in mass campaigns, or distributed in ante-natal clinics (Guyatt et al., 2002) and at schools (Yukich et al., 2020). Of immediate practical importance is the almost inevitable creation of accidental micro-mosaics of standard (pyrethroid-only) ITNs and next-generation mixture ITNs (which contain a pyrethroid and a novel insecticide). The presence of both standard ITNs and mixture ITNs in the same clusters in trials highlights this as an issue (Accrombessi et al., 2023; Mosha et al., 2022).

The paper has two aims. Firstly, to evaluate the deliberate use of micro-mosaics as an IRM strategy when compared against rotations and mixtures. The second aim is to explore the IRM implication of accidental micro-mosaics of next-generation mixture ITNs and standard (pyrethroid-only) ITNs.

## 2. Methods

### 2.1 Model Overview

We use a previously described mathematical model for the evaluation of IRM strategies (Hobbs & Hastings, 2024b) where selection in the model can be implemented as either truncation (“polytruncate”) or as a probabilistic process (“polysmooth”). In brief, the model is built in a quantitative genetics framework, assuming resistance is a polygenic trait. The model tracks the “polygenic resistance score” (PRS) of mosquito populations, which quantifies the “amount of resistance” and is measurable in bioassays (for example WHO cylinder tests). Multiple gonotrophic cycles need to be included in the model due to the hypothesised “temporal mixture” of micro-mosaics i.e., mosquitoes may encounter different insecticides throughout their lifespan. Only the “polysmooth” branch of the model can do this, so is the branch used here. Within each discrete mosquito generation, the response to selection is calculated for each insecticide in each gonotrophic cycle which is dependent on the level of resistance in the population and the efficacy of the insecticide. The key equations for the multiple gonotrophic cycle model are therefore equation 11b from (Hobbs & Hastings, 2024b), which calculates the response to selection for each gonotrophic cycle as:

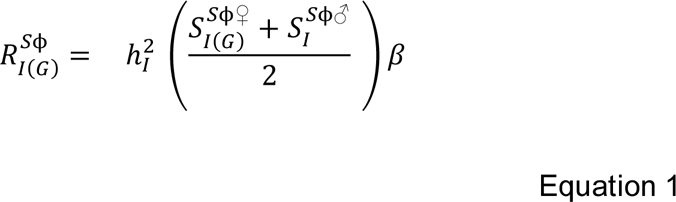

And equation 11d from (Hobbs & Hastings, 2024b) which calculates the overall response for the mosquito generation. This is achieved by weighting 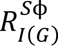 (obtained from Equation 1) by the number of females in each gonotrophic cycle:

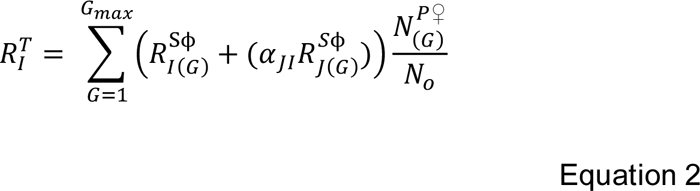

Definitions of the symbols used in the equations are in Table 1. In brief, 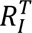 is the overall response to selection for a mosquito generation. 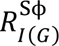 is the response to insecticide *i* for a single gonotrophic cycle. The term 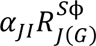 is the is the response resulting from cross resistance with insecticide *j*. The term 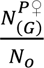 is the weighted number of females laying eggs in each gonotrophic cycle as a proportion of the total number of oviposition events across all gonotrophic cycles. A conceptual diagram of the multiple gonotrophic cycle model is given in Figure 2.

**Table 1:**
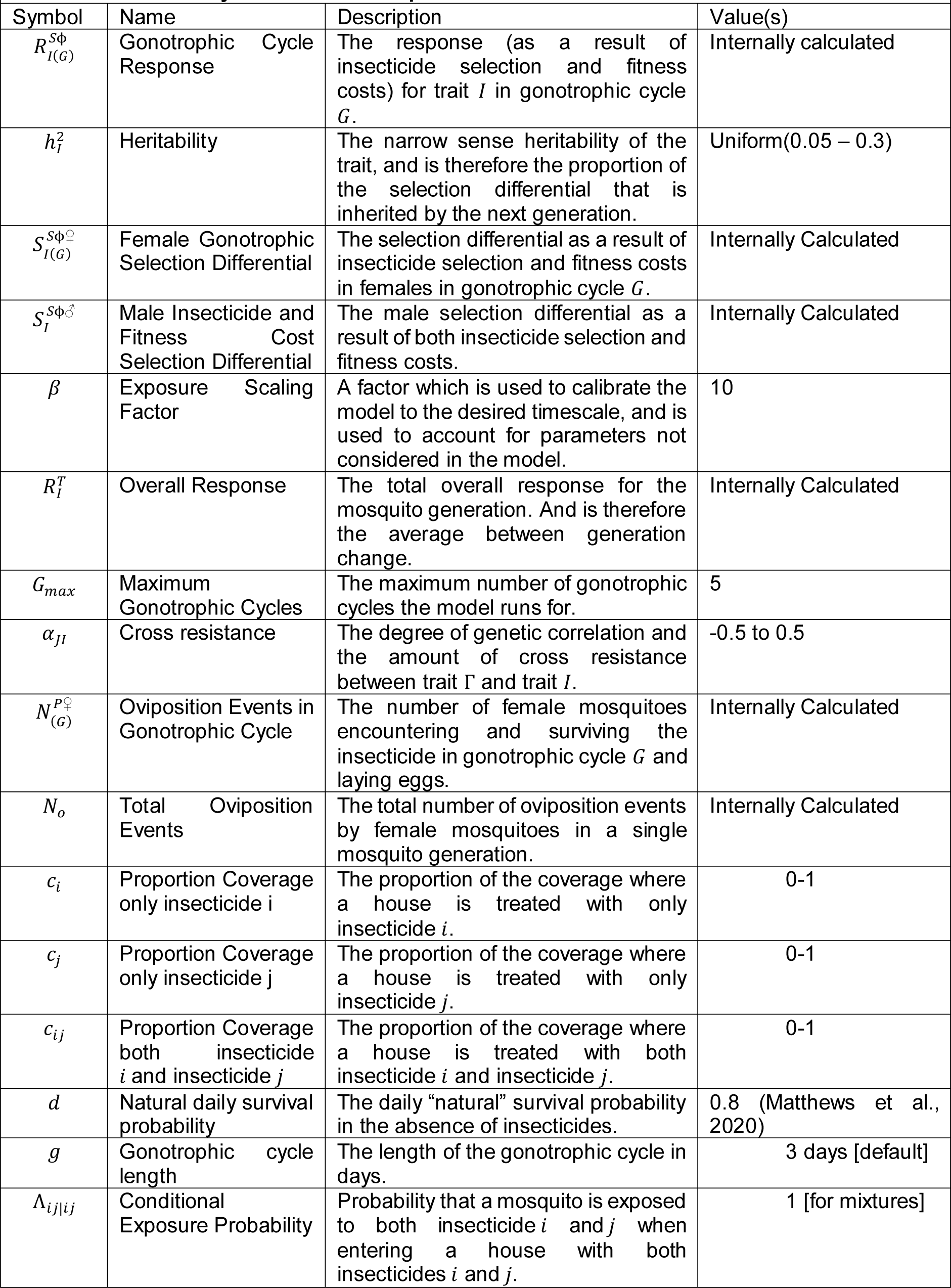
Parameter Symbols and Descriptions.

**Figure 2:**
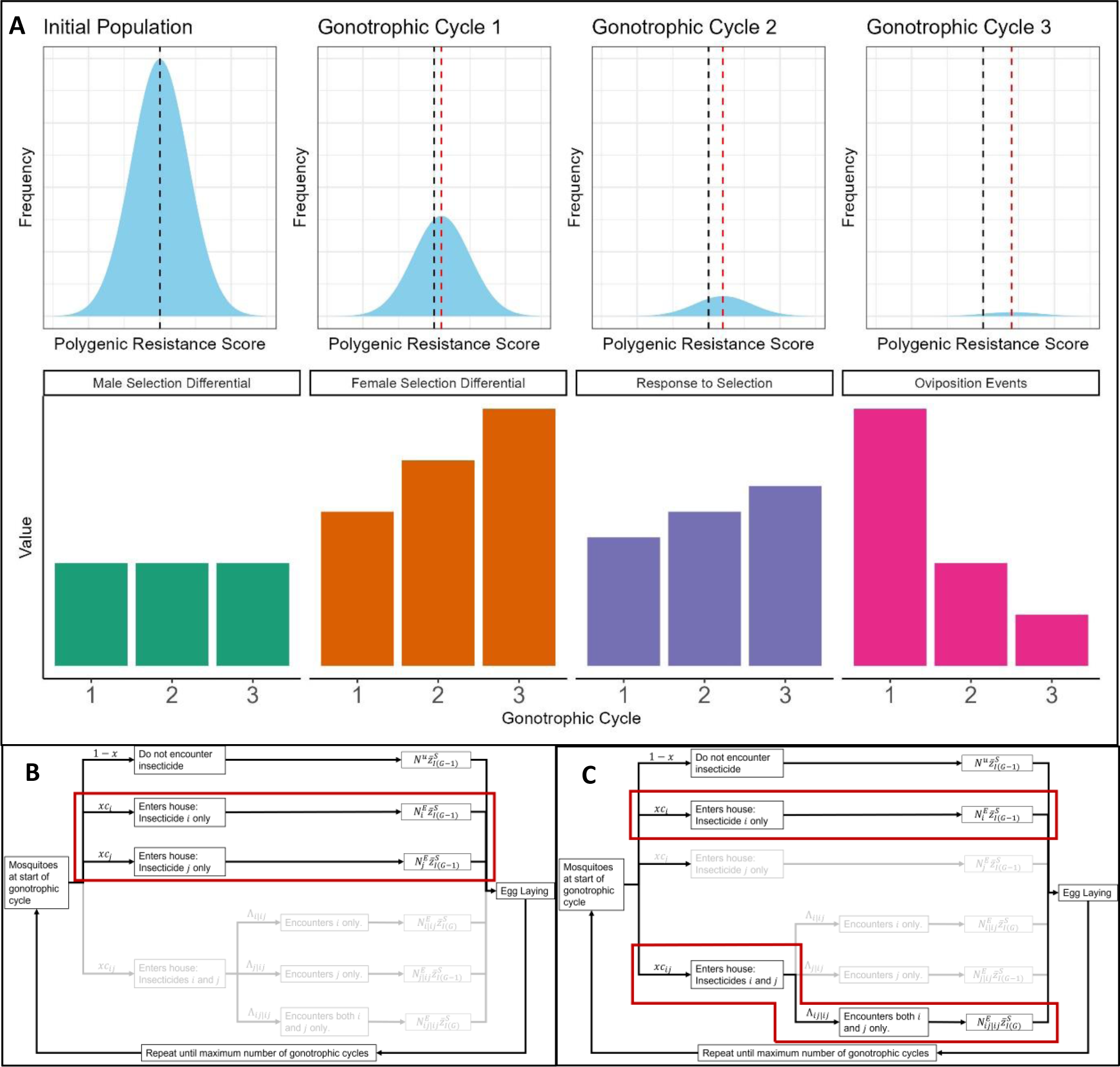
Conceptualising Multiple Gonotrophic Cycle Insecticide Selection and the Implementation of the Insecticide Resistance Management Strategies. <u>Panel A</u>: The top row of panel A shows how the contribution of each polygenic resistance score value decreases with each gonotrophic cycle because the adult mosquito population falls due to both insecticide selection and natural mortality. The average resistance after each round of selection increases (red-dashed line) because more resistant mosquitoes are more likely to survive to later gonotrophic cycles. Bottom row of panel A: The male selection differential remains constant for each gonotrophic cycle because female mosquitoes mate only once (prior to their first gonadotrophic cycles) and male mosquitoes only go through a single round of insecticide contact and selection. The female selection differential increases in each gonotrophic cycle increases in each gonotrophic cycle (difference between red and black dashed line in top row of Panel A). The response to selection increases for each gonotrophic (Equation 1), such that females laying eggs in later gonotrophic cycles will lay eggs from which more resistant individuals will hatch. However, in each gonotrophic cycle, there will be fewer females alive to lay eggs. Therefore, the response to selection for each gonotrophic is weighted by the number of oviposition events in that gonotrophic cycle (Equation 2). **<u>Panel B</u>:** Implementation of the deliberate micro-mosaics strategy, such that a proportion of households receive insecticide *i* (*c*_*i*_) and the other proportion receive insecticide *j* (*c*_*j*_). **<u>Panel C</u>:** Implementation of the accidental micro-mosaics strategy such that households have either a mixture ITN (*c*_*ij*_ and Λ_*ij*|*ij*_=1) or just a standard ITN (*c*_*i*_), where the insecticide *i* is common to both net types, as is the case with standard (pyrethroid-only) ITNs and next-generation mixture ITNs which are a mixture of a novel insecticide and a pyrethroid.

Unlike females, males do not forage for blood, so presumable have less insecticide contact, and hence undergo less selection for IR. Male Anopheline mosquitoes are frequently present in experimental hut trial collections (e.g., (Menze et al., 2020) indicating there is likely some degree of insecticide selection. However, as male mosquitoes are rarely counted or have their mortality scored in experimental hut trials the amount of insecticide selection is uncertain. As male mosquitoes do not transmit malaria their behaviour is of less public health interest than females. However, from a resistance development perspective, as males contribute 50% of the genes to the next generation they are genetically as important as females. The selection differential for males is 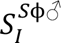 in Equation 1. Equation 1 shows this uncertainly about male selection would to be accounted for by the exposure scaling factor (β) that similarly accounts for uncertainly in heritability and female selection differential. This means uncertainty about male insecticide exposure will not affect the qualitative conclusions of the model.

The model includes multiple gonotrophic cycles, where the “natural” daily survival probability (ρ) was set at 0.8 (Matthews et al., 2020) and the length of each gonotrophic cycle (*g*) as 3 days. The maximum number of gonotrophic cycles (*G*_*max*_) used in the presented simulations is five. The exposure scaling factor (β), which is used to calibrate simulations to expected timeframes and accounts for uncertainty in the selection differentials was set at 10 as calibrated for novel insecticides (Hobbs & Hastings, 2024b).

The model allows for the inclusion of parameters which are noted to be important, yet frequently lacking from models evaluating IRM strategies, notably insecticide dosing, insecticide decay (South et al., 2020) and cross resistance (Rex Consortium, 2013). The model also allows for the evaluation of multiple different IRM strategies in the same framework allowing for direct comparisons between IRM strategies.

The model code is written in R, with statistical analysis and data visualisation also conducted in R (R Core Team, 2020).

### 2.2. Scenario Overviews: Description of Insecticide Resistance Management Strategies

Simulations in this study are separated into two distinct categories. Scenarios 1A, 1B, 1C, 1D investigate the deliberate use of micro-mosaics as an IRM strategy. Scenario 2 investigates the accidental occurrence of micro-mosaics as may be expected due to multiple distribution channels, with a specific focus on deployment of the new generation of mixture ITNs onto an existing background of standard (pyrethroid-only) ITNs.

In Scenario 1 micro-mosaics are evaluated as a deliberately implemented IRM strategy and are compared against rotations, full-dose mixtures, and half-dose mixtures in four sub-scenarios:

- Scenario 1A: quantifies the performance of perfect 50:50 deliberate micro-mosaics for novel insecticides.
- Scenario 1B: quantifies the performance of perfect 50:50 deliberate micro-mosaics when there is already pre-existing resistance in the mosquito population.
- Scenario 1C: quantifies the performance of deliberate micro-mosaics under increased biological and operational realism.
- Scenario 1D: quantifies the performance of deliberate micro-mosaics in the presence of insecticide decay.

Conceptually, 50:50 deliberate micro-mosaics, rotations and half-dose mixtures can be viewed as halving the amount of each insecticide deployed in different ways:

- For a 50:50 deliberate micro-mosaic (*c*_*i*_= 0.5, *c*_*j*_ = 0.5, Figure 2 Left Panel), the amount of each insecticide is halved for each deployment. With 50% of households receiving insecticide *i* and 50% of households receiving insecticide *j*.
- For rotations the selection by each insecticide can be considered halved over time, where all households receive insecticide *i* in one deployment (*c*_*i*_= 1), and then all households receive insecticide *j* in the next deployment (*c*_*j*_ = 1).
- For half-dose mixtures the amount of insecticide is halved by dose. Where all households receive both insecticides (*c*_*ij*_ = 1) every deployment, but the dose of these insecticides is halved.

For each strategy (deliberate micro-mosaics, rotations, half-dose mixtures and full-dose mixtures, Figure 1), the same 5000 biological parameter sets were used for each strategy within a Scenario allowing for direct comparisons between strategies. Latin hypercube sampling (Carnell, 2020) of uniform distributions were used to generate the parameter sets for each scenario The exact parameter sets may vary between scenarios, as detailed later.

For Scenario 1 the simulation rules are as follows. First, the withdrawal threshold (the recommended point at which insecticides should no longer be used) (Table 1) was set. At this point the insecticide is designated as having “failed” and is therefore withdrawn and cannot be deployed at the next deployment interval. For Scenarios 1A, 1C and 1D the withdrawal threshold was set at 10% bioassay survival (WHO, 2018). For Scenario 1B (where insecticides start at 10% bioassay survival) the withdrawal threshold was set at 20% bioassay survival. The choice of bioassay survival for a “failed insecticide” is essentially an arbitrary cut-off (both practically and in simulations) which is not informed by any threshold at which vector control fails. In models assuming a monogenic basis of resistance the threshold for withdrawal/failure is usually set at a 50% resistance allele frequency regardless of the biological system (Birget & Koella, 2015; Caprio & Tabashnik, 1992; Levick et al., 2017; Roush, 1998; Slater et al., 2017; Sudo et al., 2018) although a 20% resistance allele frequency has also been used (Argentine et al., 1994). Questions therefore arise whether this choice of threshold also impacts the comparative performance of IRM strategies.

Simulations are terminated when either the 500-generation (∼50 years, assuming 10 generations per year) limit is reached or when the strategy has failed. A failed strategy occurs for mixtures and deliberate micro-mosaic deployments when either of the two insecticides is withdrawn, as the strategy is no longer able to be implemented. For rotations the simulation terminates if no insecticide is available to be rotated to, that is the other insecticide has already “failed”, this is when the rotation strategy can no longer be implemented. The duration of the simulation in years (assuming 10 generations per year) is defined as the “strategy lifespan”.

The deployment interval was 10 generations for Scenarios 1A, 1B and 1C because this allows for a high resolution of strategy lifespan. Extending the deployment interval to 30 generation (∼ 3 years as for ITNs) decreases the resolution, such that any benefits between two strategies are less clearly seen. i.e., if insecticides “fail” 2 generations into a 10-generation interval there is only 8 more generations until the simulation terminates, whereas if insecticides “fail” 2 generations into a 30-generation deployment interval then there are still 28 generations until the simulation terminates and the strategy is identified as having reached its strategy lifespan. In models not including insecticide decay the qualitative conclusions between 10 and 30 generation deployment intervals are similar (N. Hobbs et al., 2023). Considering, half-dose mixtures, rotations, and deliberate micro-mosaics have the same “total amount of insecticide”, this higher resolution is required to determine if there is any difference in strategy lifespans. As Scenario 1D includes insecticide decay, the deployment interval was changed to 30 generations (∼3 years).

The primary outcome was the duration of the simulation (“strategy lifespan”). As our main interest is in the performance of deliberate micro-mosaics, we compare the “strategy lifespan” for deliberate micro-mosaics against each other strategy (rotations, full-dose mixtures and half-dose mixtures) and define the deliberate micro-mosaic to have won, drawn, or lost. The performance of rotations, half-dose mixtures, full-dose mixtures against each other has been evaluated elsewhere (N. Hobbs et al., 2023). If two strategies have the same “strategy lifespan” for a parameter set, they are compared on the secondary outcomes of the combined peak bioassay survival (insecticide *i* + insecticide *j*) reached in the simulation as a measure of the “total amount of resistance”, where the strategy with the lower “total amount of resistance” therefore performing better from an IRM perspective.

For the primary analysis we present the distributions of the differences in the “strategy lifespans” between deliberate micro-mosaics and each of the comparator strategies (rotations, half-dose mixtures, full-dose mixtures) as histograms. Where strategies draw, the distributions of the differences in the combined peak bioassay survival reached in the simulations are presented as histograms.

Sensitivity analysis was conducted using generalised additive models (GAMs). The exact details of the GAMs are detailed later. GAMs allow for the impact of the parameter on the outcome to be seen with any non-linear relationships captured to give an understanding of areas of the parameter space where different strategies perform better or worse.

### 2.3 Scenario 1A: Evaluating Deliberate Micro-Mosaics with Novel Insecticides

Scenario 1A evaluates the use of deliberate micro-mosaics assuming the insecticides are novel (initial starting bioassay survival is 0%) and is summarised in Table 2. Simulations were run with cross resistance (α_*IJ*_= -0.5, -0.3, -0.1, 0, 0.1, 0.3, 0.5) and therefore each strategy was therefore run for a total of 35,000 simulations. The model was run assuming the standard deviation (σ_*I*_) remained fixed throughout the simulations (σ_*I*_ = 50).

**Table 2:** Design Summary for Scenario 1A.

| Table 2: Design Summary for Scenario 1A |  |  |  |  |  |
| --- | --- | --- | --- | --- | --- |
| IRM Strategy | | Parameter [5000 sets] | | Cross Resistance, $\alpha_{IJ}$ | Outcomes |
| Micro-mosaics | X | Heritability, $h_I^2$ (0.05 – 0.3) | X | -0.5 | Primary:<br>Simulation Duration<br>("Strategy Lifespan")<br>Secondary:<br>Peak Bioassay Survival |
| | | Male Fitness Cost, $S_I^{\phi\sigma}$ (0.04 – 0.58) | | -0.3 | |
| Rotations | | Female Fitness Cost, $S_I^{\phi\varphi}$ (0.04 – 0.58) | | -0.1 | |
| Half-Dose Mixture | | Male Exposure, $m$ (0-1) as a proportion of female exposure. | | 0 | |
| | | Female Insecticide Exposure, $x$ (0.4 – 0.9) | | 0.1 | |
| Full -Dose Mixture | | Intervention Coverage, $C$ (0.5 – 0.9) | | 0.3 | |
| | | Dispersal, $\theta$ (0.1 – 0.9) | | 0.5 | |

Sensitivity analysis was conducted using generalised additive models (GAMs) each randomly sampled parameter (Heritability, Female Insecticide Exposure, Male Insecticide Exposure, Female Fitness Costs, Male Fitness Costs, Intervention Coverage and Dispersal) against the difference in the strategy lifespan between the deliberate micro-mosaics simulations and the comparator simulations. Due to the identification from the primary analysis of the importance of cross resistance, models were fit were fit separately for negative cross resistance, no cross resistance and positive cross resistance.

### 2.4 Scenario 1B: Evaluating Deliberate Micro-Mosaics under Higher Resistance Levels

Scenario 1B evaluates the use of deliberate micro-mosaics assuming the insecticides have pre-existing resistance (initial starting bioassay survival is 10%) and is summarised in Table 3. As simulations were run at higher resistance levels, the standard deviation was allowed to vary with the magnitude of the mean PRS and the fitness costs were allowed to vary with the standard deviation (Hobbs & Hastings, 2024b), keeping the effect constant. Simulations were run including cross resistance (-0.5, -0.3, -0.1, 0, 0.1, 0.3 and 0.5) and therefore each strategy was therefore run for a total of 35,000 simulations. Sensitivity analysis using GAMs was conducted as was described for the Scenario 1A sensitivity analysis.

**Table 3:**
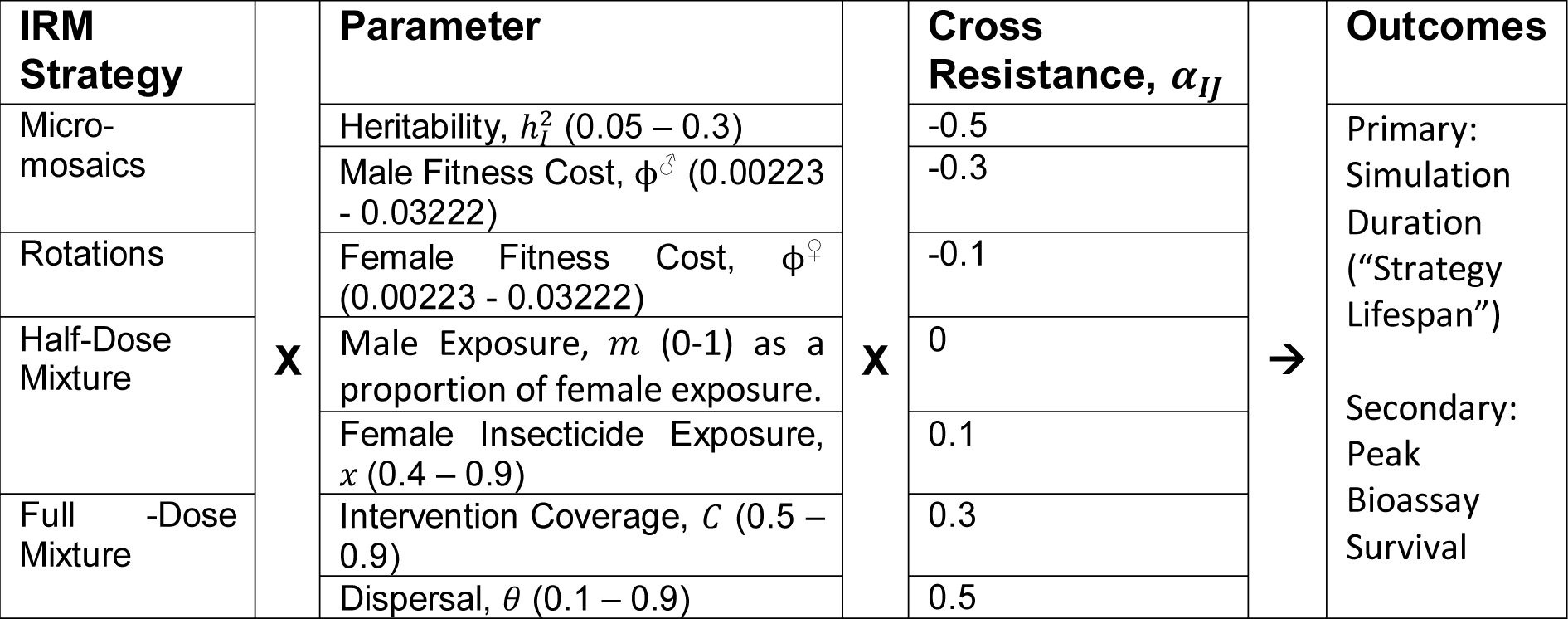
Design Summary for Scenario 1B.

### 2.5. Scenario 1C: Evaluating Deliberate Micro-Mosaics with Biological and Operational Realism

Scenario 1C evaluates the use of deliberate micro-mosaics under additional biological/operational realism. This is to ensure that any benefits (or not) of deliberate micro-mosaics are not restricted to “perfect” deployments, which in real-world settings would be unlikely. This is especially important as for deliberate micro-mosaics it would be unrealistic to expect a perfect 50:50 ratio of both insecticides and, therefore, the rate of evolution to either insecticide would no longer be expected to be at the same rate. There are a few reasons why an imperfect 50:50 ratio may occur. First is supply chain/delivery issues, second is the economic cost of each insecticide, third is deployment issues. It is therefore important to check the implications of imperfect deliberate micro-mosaic deployments. The proportion coverage of only insecticide *i* (*c*_*i*_) was randomly sampled between 0.4 and 0.6, with the corresponding proportion coverage only insecticide *j* (*c*_*j*_) being 1 − *c*_*i*_ . Note that *c*_*i*_ and *c*_*j*_ are fixed throughout the simulations. In real-world scenarios it would be expected for coverages to vary in each deployment. The insecticides were also allowed to have unique properties (e.g., heritability and starting resistance). For each strategy, the same 40000 biological parameter sets were used (Table 4). The model was run assuming the standard deviation (σ_*I*_) remained fixed throughout the simulations (σ_*I*_ = 50). All simulations are capped at 500 generations (∼50 years).

**Table 4:** Design Summary for Scenario 1C.

| IRM Strategy |  | Parameter [40000 sets] |  | Outcomes |
| --- | --- | --- | --- | --- |
| Micro-mosaics | X | Heritability, $h_i^2$ and $h_j^2$ (0.05 – 0.3) | → | Primary:<br>Simulation Duration<br>("Strategy Lifespan")<br>Secondary:<br>Total End<br>Bioassay<br>Survival |
| | | Male Fitness Cost, $S_i^{\phi\sigma}$ and $S_j^{\phi\sigma}$ (0.04 – 0.58) | | |
| Rotations | | Female Fitness Cost, $S_i^{\phi\sigma}$ and $S_j^{\phi\sigma}$ (0.04 – 0.58) | | |
| | | Male Exposure, $m$ (0-1) as a proportion of female exposure. | | |
| | | Female Insecticide Exposure, $x$ (0.4 – 0.9) | | |
| Half-Dose Mixture | | Intervention Coverage, $C$ (0.5 – 0.9) | | |
| | | Dispersal, $\theta$ (0.1 – 0.9) | | |
| | | Cross Resistance, $\alpha_{IJ}$ (-0.5 – 0.5) | | |
| Full -Dose Mixture | | Starting Polygenic Resistance Score, $\bar{z}_i$ and $\bar{z}_j$ (0 – 80) | | |
| | | For Micro-Mosaics Only:<br>Coverage Insecticide $i$, $c_i$ (0.4 – 0.6)<br>Coverage Insecticide $j$, $c_j = 1 - c_i$ | | |

Sensitivity analysis was conducted using GAMs fitting over each randomly sampled parameter against the difference in the strategy lifespan between the deliberate micro-mosaics simulations and the comparator simulations. These models were fit separately for negative cross resistance, no cross resistance and positive cross resistance.

### 2.6 Scenario 1D: Evaluating Deliberate Micro-Mosaics with Insecticide Decay

Scenario 1C evaluates the use of deliberate micro-mosaics in the presence of insecticide decay. Insecticide decay has been frequently highlighted as an important consideration for mixtures (Curtis, 1985) and as micro-mosaics are intended to work as “temporal mixtures” insecticide decay may be an issue here too.

The insecticides remain having the same parameter values as each other (initial bioassay survival, heritability, fitness costs). However, the two insecticides are allowed to decay at different rates. For insecticide *i* the base decay rate was set as 0.005 (“slow”), 0.015 (“default”) and 0.025 (fast); and the base decay rate for insecticide *j* was fixed at 0.015 throughout. We can therefore compare the implication of the two insecticides decaying slower, equally, or faster to one another, without the additional noise of the insecticides having other different properties (such as heritability and fitness costs). The threshold generation was 20 (∼2 years), after which both insecticides have an increased decay rate of 0.08, indicating the net material has been sufficiently damaged to accelerate the rate of decay. The deployment interval was set at 30 generations (∼3 years), and therefore this mimics the deployment of insecticides as ITNs.

For each strategy, the same 5000 biological parameter sets were used (Table 5), Simulations were run including cross resistance (-0.3, 0 and 0.3) and therefore each strategy was run for a total of 15,000 simulations for each decay profile permutation. The model was run assuming the standard deviation (σ_*I*_) remained fixed throughout the simulations (σ_*I*_ = 50).

**Table 5:**
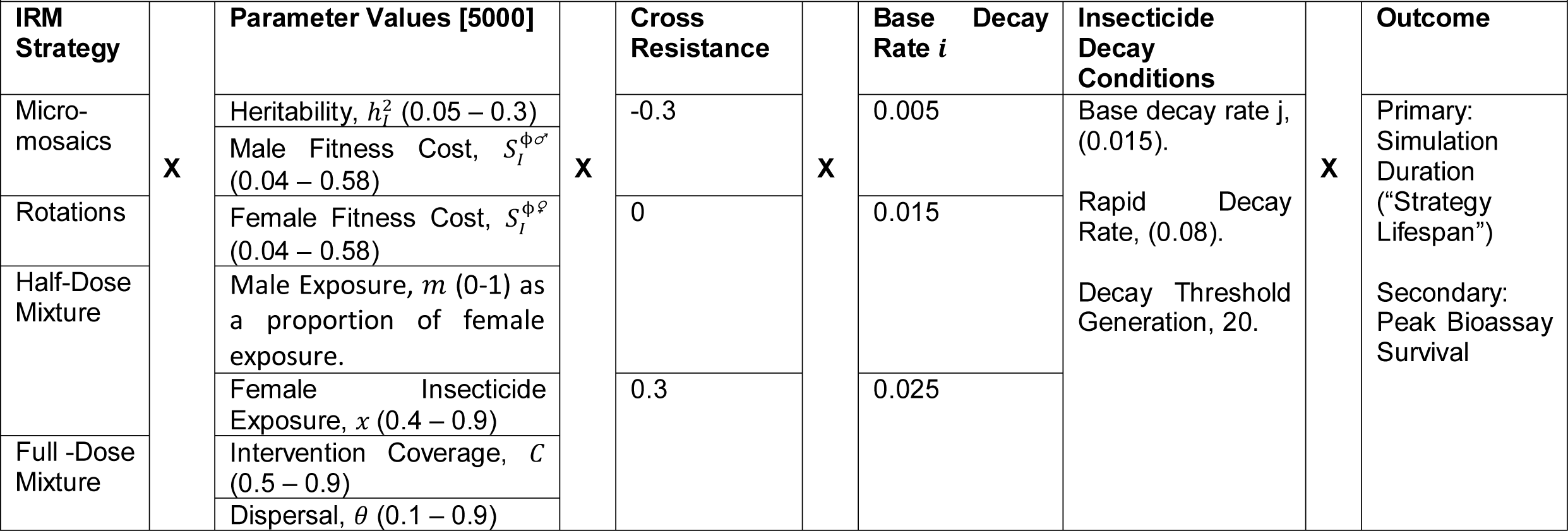
Design Summary for Scenario 1D.

Sensitivity analysis was conducted using GAMs fitting over each randomly sampled parameter against the difference in the strategy lifespan between the deliberate micro-mosaics simulations and the comparator simulations. These models were fit separately for negative cross resistance, no cross resistance and positive cross resistance and for each decay profile permutation.

### 2.7. Scenario 2: Implications of Accidental Micro-Mosaics of Standard and Mixture ITNs

Scenario 2 investigates co-deployment of both next-generation ITNs (which contain a pyrethroid insecticide and a novel insecticide) and standard (pyrethroid-only) ITNs in the same village; as identified as occurring during cluster-RCTs (Accrombessi et al., 2023; Mosha et al., 2022). We term this “accidental micro-mosaics” (Figure 1). Accidental micro-mosaics simulations were implemented using the combinations capability of the model (Figure 2, right panel), which allows for houses to be given either insecticide *i* (with proportional coverage *c*_*i*_) or insecticides *i* and *j* (proportional coverage *c*_*ij*_), with proportional coverage *c*_*j*_ set as 0. The coverage of *c*_*ij*_ is turned into a mixture by setting the conditional encounter rate (Λ_*ij*|*ij*_) to 1 for both males and females, such that mosquitoes are guaranteed to encounter both insecticides simultaneously. For all simulations *c*_*ij*_ + *c*_*i*_ = 1. The impact of mixture coverage within an accidental micro-mosaic (*c*_*ij*_ , set as 0.25, 0.5, 0.75) can therefore be explored against (i) the standard (pyrethroid-only) ITN only simulations (expected lower benchmark) where the proportion of the coverage that is the standard ITN is *c*_*i*_ = 1 and (ii) mixture only ITNs simulations (expected upper benchmark) where the proportional coverage of the mixture is *c*_*ij*_ = 1.

Mixtures were allowed to either be full-dose or half-dose (retaining 50% efficacy). This means for the accidental micro-mosaics with mixtures at half-dose, the pyrethroid-only ITNs are at full-dose and the insecticides in the mixture are each at half-dose. To enable these scenarios to be run, the underlying model code was adapted to allow for the deployment of standard ITN (full-dose), and mixture ITN (with both insecticides at half-dose). These code modifications however prevent the inclusion of insecticide decay in the simulations. Nevertheless, the results will still provide useful information regarding “accidental” micro-mosaic deployments, a real-world issue which has not been explicitly evaluated previous. Note, if half-dose mixtures are used in the accidental micro-mosaic simulations, the comparator upper benchmark simulations use half-dose mixture ITNs also.

As there is likely to already be high levels of IR to the pyrethroid where next-generation ITNs are used, the pre-existing IR level to the pyrethroid was varied (0.5, 10, 20, 50, 80% bioassay survival). For each coverage-resistance scenario, the same 5000 biological parameter sets were used (Table 6), allowing for direct comparisons between the accidental micro-mosaic simulations, mixture only simulations and pyrethroid only simulations.

**Table 6:** Design Summary for Scenario 2.

| Table 6: Design Summary for Scenario 2 |  |  |  |  |  |  |  |  |  |  |  |  |
| --- | --- | --- | --- | --- | --- | --- | --- | --- | --- | --- | --- | --- |
| Coverage standard ITN ($c_i$) | Coverage mixture ITN ($c_{ij}$) | | Start Pyrethroid Bioassay Survival (%) | | Dosing of the Mixture | | Parameter | | Outcomes | | Evaluation | |
| 0 | 1 [Upper Benchmark] | X | 0.1 | X | Full-Dose (100% efficacy) | X | Heritability, $h_l^2$ (0.05 – 0.3) | → | Bioassay survival change insecticide i.<br>Bioassay survival change insecticide j.<br>Total (combined) bioassay change. | → | Micro-mosaic deployments are compared against:<br>Upper benchmark of mixture ITN only.<br>Lower benchmark of standard ITN only. | |
| 0.25 | 0.75 | | 10 | | | | Male Exposure, $m$ (0-1) as a proportion of female exposure. | | | | | |
| 0.5 | 0.5 | | 20 | | | | Female Exposure, $x$ (0.4 – 0.9) | | | | | |
| 0.75 | 0.25 | | 50 | | Half-Dose (retaining 50% efficacy) | | Intervention Coverage, $c$ (0.5 – 0.9) | | | | | |
| 1 [Lower Benchmark] | 0 | | 80 | | | | Dispersal, $\theta$ (0.1 – 0.9) | | | | | |

In Scenario 2, simulations were run for 200 generations (∼20 years) with continuous deployment and at the end of the simulations the end bioassay survival for each insecticide was extracted. A time horizon of 20 years was used as after this the expectation is standard “pyrethroid” only ITNs will no longer be used.

The differences between the end bioassay survival for accidental micro-mosaic simulations and the corresponding benchmark simulations were calculated for each insecticide. The lower benchmark simulations were standard (pyrethroid-only) ITNs only and the upper benchmark simulations were mixture ITNs only.

For the pyrethroid insecticide this was calculated as:

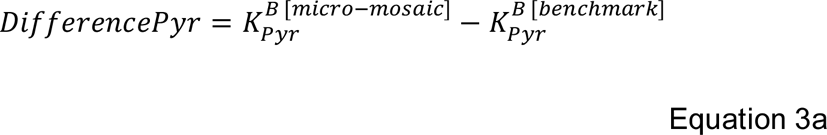

Where the value 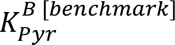 is the end bioassay survival (after 200 generations) for the pyrethroid insecticide from the benchmark simulations, being either standard (pyrethroid-only) ITNs only (lower benchmark) or mixture ITNs only (upper benchmark). The value 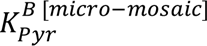 is the end bioassay survival (after 200 generations) for the pyrethroid insecticide from the accidental micro-mosaic simulations. The difference for the novel insecticide is also calculated:

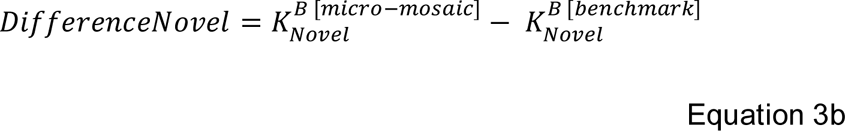

The value 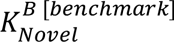 is the end bioassay survival (after 200 generations) from the benchmark simulations for the novel insecticide. The value 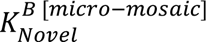 is the end bioassay survival (after 200 generations) for the novel insecticide from the accidental micro-mosaic simulations. The overall difference in the end bioassay survivals is then calculated:

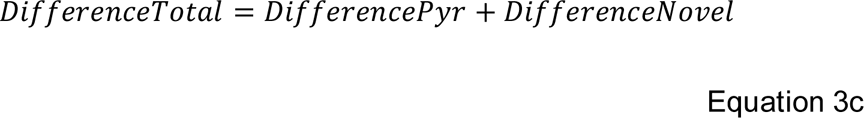

The primary analysis examined these differences in IR level (*DifferencePyr*, *DifferenceNoval*, *DifferenceTotal*) between the accidental micro-mosaic simulations and the corresponding expected lower benchmark of pyrethroid-only ITNs and the expected upper benchmark of the mixture-only ITNs. These were reported as histograms.

Regression classification trees were used to identify the parameter permutations where accidental micro-mosaics outperformed mixture ITNs only (the upper benchmark) or were outperformed by standard (pyrethroid-only) ITNs only (the lower benchmark). Regression classification trees were fit using rpart (Therneau & Atkinson, 2019) with a minimum allowable end node size of 50 and maximum depth of five and the regression classification trees were plotted using rpart.plot (Milborrow, 2020). The outcomes were classed as accidental micro-mosaics win, accidental micro-mosaics lose or the strategies draw based on the & *DifferenceTotal*. The predictor variables were female insecticide exposure, male insecticide exposure, heritability, intervention coverage, dispersal, initial pyrethroid resistance and the proportion of coverage mixture (*c*_*ij*_). Four regression classification trees were fit to allow for comparison against both the upper and lower benchmarks when assuming both half-dose and full-dose mixtures. Each regression classification tree was fit against a randomly sampled 70% of the simulations. The accuracy of the fitted classification tree was then tested against the remaining 30% of simulations.

## 3. Results

### 3.1. Scenario 1A: Evaluating Deliberate Micro-Mosaics with Novel Insecticides under Optimal Conditions

When comparing deliberate micro-mosaics against rotations for novel insecticides (Figure 3, Top Row), deliberate micro-mosaics outperform rotations by either having a longer strategy lifespan or maintaining lower levels of resistance when there is either no cross resistance or negative cross resistance. However, when there is positive cross resistance, rotations generally outperform deliberate micro-mosaics. The differences in strategy lifespans between deliberate micro-mosaics and rotations is often small, and the benefit of either strategy over the other is typically less than 5 years (Figure 3, Top Row). A total of 6680 (13.36%) simulations had deliberate micro-mosaics lasting 5 years or more than rotations. While a total of 2087 (4.17%) simulations had rotations lasting 5 years or more than micro-mosaics. Sensitivity analysis highlights the comparative advantage of deliberate micro-mosaics over rotations when there is negative cross resistance and when heritability, female insecticide exposure and intervention coverage are high (Figure 4).

**Figure 3:**
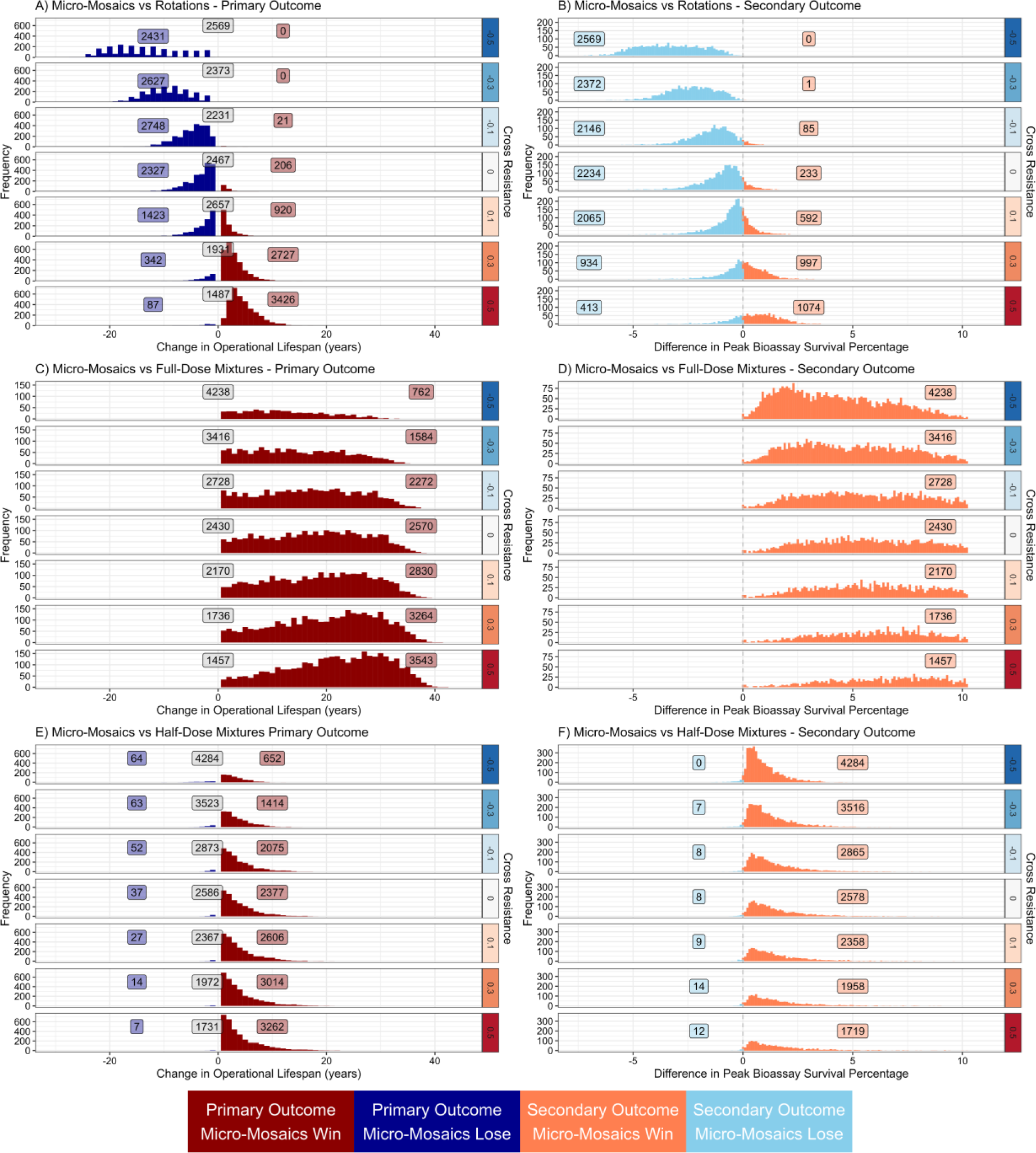
Scenario 1A - Comparing Micro-Mosaics versus Rotations (top), Full-Dose Mixtures (middle) and Half-Dose Mixtures (bottom) for novel insecticides. Plots on the left evaluate the primary outcome i.e., the difference in strategy lifespan. Plots on the right evaluate. Values to the left (dark blue) indicate the micro-mosaics strategy had a longer strategy lifespan, and values to the right (dark red) indicate the comparator strategy had a longer strategy lifespan. Each plot is stratified by the degree of cross resistance between the two insecticides. The numbers in each plot panel indicate the total of wins, losses and draws, with the grey boxes the number of draws in the strategy lifespan. Draws are evaluated on the secondary outcome of the peak bioassay survival shown in the plots on the right. Values to the left (light blue) indicate the micro-mosaics strategy had a lower peak bioassay survival, and values to the right (light red) indicate the comparator strategy had a lower peak bioassay survival.

**Figure 4:**
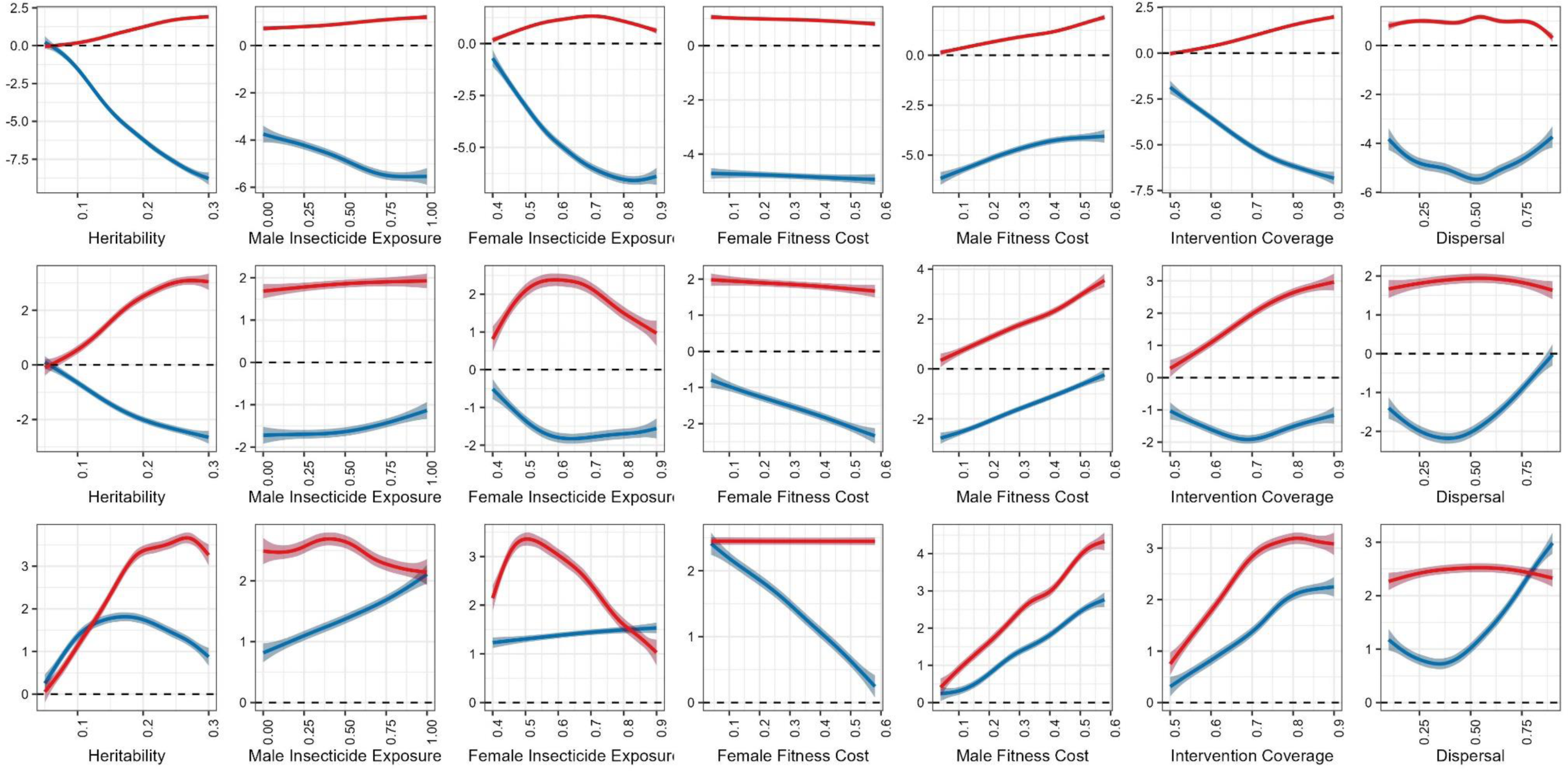
Generalised Additive Models for Scenario 1A. Top row is for negative cross resistance (α_*ij*_ = -0.1, -0.3 and -0.5), middle row is no cross resistance (α_*ij*_ = 0), and bottom row is for positive cross resistance (α_*ij*_ = 0.1, 0.3 and 0.5). The y axis is the difference in Strategy Lifespan (in years) between the micro-mosaics strategy and the corresponding comparator strategies of rotations (blue) and half-dose mixtures (red). The full-dose mixture strategy was not included due to outperforming the micro-mosaics simulations for all simulations. The dashed line at 0 indicates the micro-mosaic and comparator strategy performed equally, and values above this line indicate the comparator strategy had a longer strategy lifespan, and values below the line indicate the micro-mosaic strategy had a longer strategy lifespan.

When comparing deliberate micro-mosaics against half-dose mixtures for novel insecticides (Figure 3, middle row), regardless of the level of cross resistance, half-dose mixtures outperform deliberate micro-mosaics by having either a longer strategy lifespan, or in comparisons where the strategy lifespans were equal ending the simulations at lower levels of resistance. However, the differences in strategy lifespans were typically short being less than 5 years (Figure 3, middle row), with only 4827 (9.65%) total simulations extending the lifespan 5 years or more.

When comparing deliberate micro-mosaics against full-dose mixtures for novel insecticides (Figure 3, bottom row), it can be seen, regardless of the level of cross resistance, that full-dose mixtures outperform deliberate micro-mosaics. When the full-dose mixture strategy had a longer strategy lifespan, this was typically more than 5 years (Figure 3, bottom row). In total 15359 (30.72%) of simulations ended with full-dose mixtures having a strategy lifespan of 5 or more years than deliberate micro-mosaics.

### 3.2. Scenario 1B: Evaluating Deliberate Micro-Mosaics under Higher Resistance Levels

Comparing deliberate micro-mosaics against rotations in simulations starting at higher IR levels (Figure 5, top row), deliberate micro-mosaics generally outperform rotations when there is negative cross resistance. However, when there is no cross resistance there is a shift to rotations generally outperforming deliberate micro-mosaics. When there is positive cross resistance, rotations consistently outperform deliberate micro-mosaics. When the no-cross-resistance simulations are compared against when using novel insecticides (Scenario 1A, Figure 3, top row), the general qualitative conclusions of these simulations flip with deliberate micro-mosaics going from performing generally better than rotations when both insecticides are novel to performing generally worse than rotations when the simulations are run at higher resistance levels (Scenario 1B: higher IR levels). Nevertheless, the quantitative differences in strategy lifespans remain small (especially when compared against full-dose mixtures), typically being less than 5 years. A total of 7059 (14.12%) simulations had deliberate micro-mosaics lasting 5 years or more than rotations. While a total of 5069 (4.17%) simulations had rotations lasting 5 years or more than deliberate micro-mosaics. The sensitivity analysis (Figure 6), highlights that the benefit of deliberate micro-mosaics over rotations under higher resistance scenarios is increased when male fitness costs increase or female insecticide exposure increases. And that the benefit of deliberate micro-mosaics over rotations decreases when female fitness costs increase or the intervention coverage increases.

**Figure 5:**
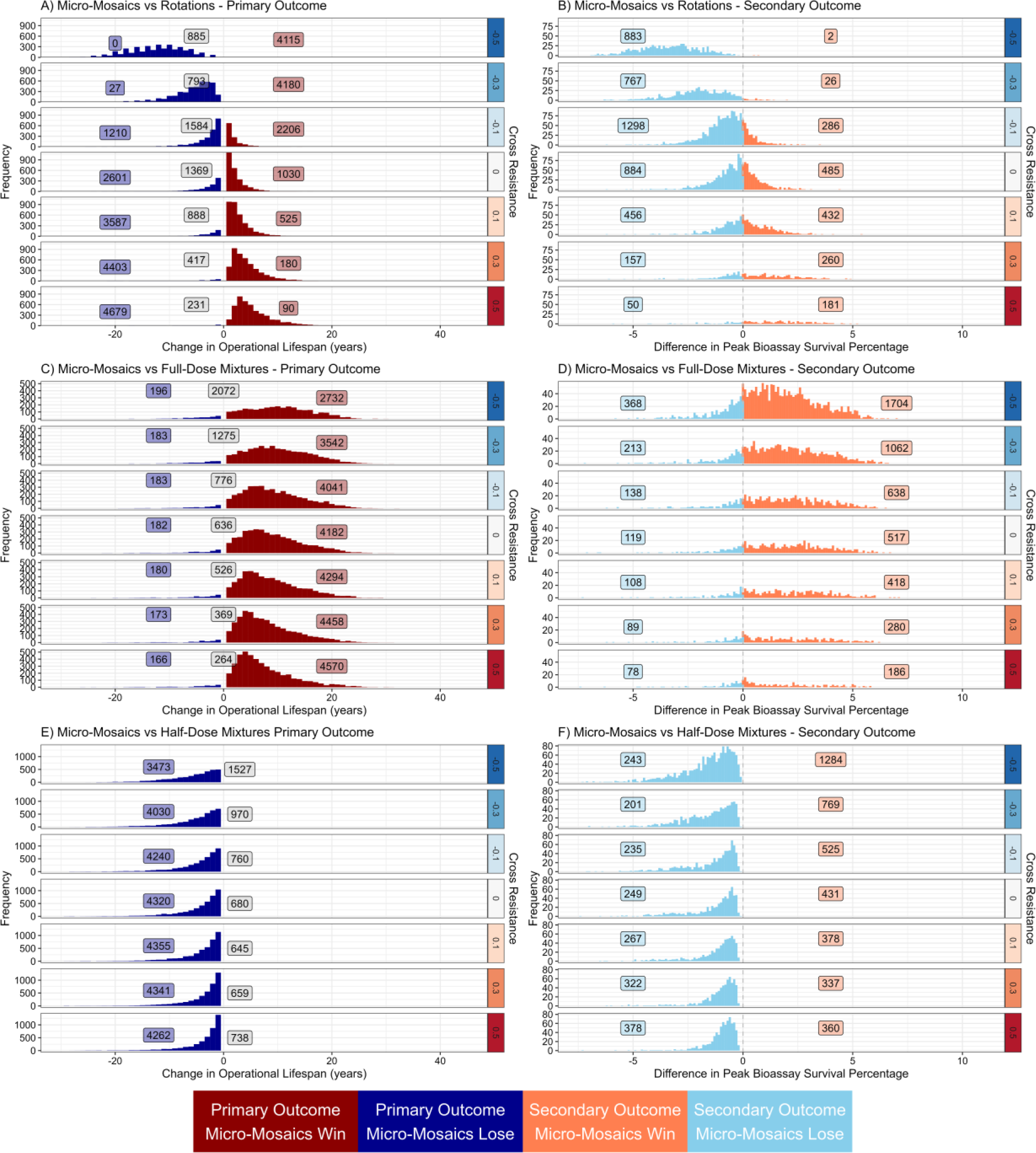
Scenario 1B - Comparing Micro-Mosaics versus Rotations (top), Full-Dose Mixtures (middle) and Half-Dose Mixtures (bottom) at higher levels of resistance. Plots on the left evaluate on the primary outcome of the difference in strategy lifespan. Values to the left (dark blue) indicate the micro-mosaics strategy had a longer strategy lifespan, and values to the right (dark red) indicate the comparator strategy had a longer strategy lifespan. Each plot is stratified by the degree of cross resistance between the two insecticides. The numbers in each plot panel indicate the total of wins, losses and draws, with the grey boxes the number of draws in the strategy lifespan. Draws are evaluated on the secondary outcome of the peak bioassay survival shown in the plots on the right. Values to the left (light blue) indicate the micro-mosaics strategy had a lower peak bioassay survival, and values to the right (light red) indicate the comparator strategy had a lower peak bioassay survival.

**Figure 6:**
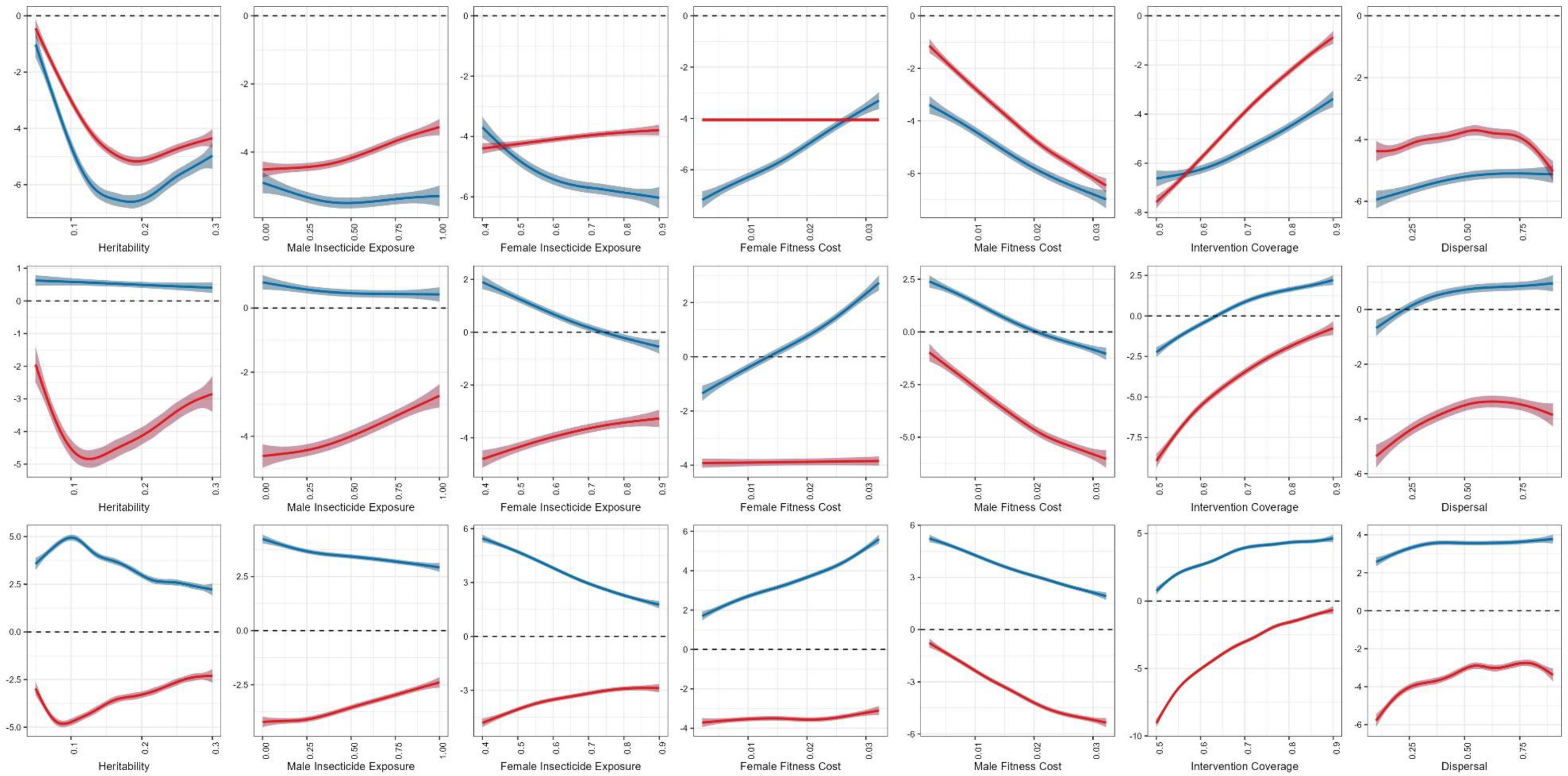
Generalised Additive Models for Scenario 1B. Top row is for negative cross resistance (α_*ij*_ = -0.1, -0.3 and -0.5), middle row is no cross resistance (α_*ij*_ = 0), and bottom row is for positive cross resistance (α_*ij*_ = 0.1, 0.3 and 0.5). The y axis is the difference in Strategy Lifespan (in years) between the micro-mosaics strategy and the corresponding comparator strategies of rotations (blue) and half-dose mixtures (red). The full-dose mixture strategy was not included due to outperforming the micro-mosaics simulations for all simulations. The dashed line at 0 indicates the micro-mosaic and comparator strategy performed equally, and values above this line indicate the comparator strategy had a longer strategy lifespan, and values below the line indicate the micro-mosaic strategy had a longer strategy lifespan.

When comparing deliberate micro-mosaics against half-dose mixtures in simulations run at higher IR levels (Figure 5, middle row), deliberate micro-mosaics outperform the half-dose mixtures regardless of the level of cross resistance. This is a qualitatively different result compared to when evaluating using novel insecticides (Scenario 1A, Figure 3, middle row) where half-dose mixtures were found to outperform deliberate micro-mosaics regardless of the amount of cross resistance. Nevertheless, the quantitative difference in strategy longevities is still typically less than 5 years. This does highlights the issue of the need to consider improving how strategies are evaluated with regards to failure thresholds, whereby different strategies may perform better or worse at different resistance levels.

When comparing deliberate micro-mosaics against full-dose mixtures in simulations run at higher IR levels (Figure 5, bottom), it can be seen that full-dose mixtures outperform deliberate micro-mosaics regardless of the level of cross resistance. However, unlike the novel insecticide simulations (Scenario 1A, Figure 3, bottom row), there were very infrequent occasions (less than 4% of simulations) where the deliberate micro-mosaic strategy was found to outperform the full-dose mixture strategy. In general, full-dose mixtures had a strategy lifespan 5 to 15 years longer than micro-mosaics (Figure 5, bottom), although this is notably shorter than the 10-30 years when both insecticides are novel.

### 3.3. Scenario 1C: Evaluating Deliberate Micro-Mosaics Under Operational and Biological Realism

Scenario 1C (Table 3) extended the simulations to allow for the insecticides to have unique properties (e.g., heritability, fitness costs) and for the insecticides deployed as micro-mosaics to be deployed at unequal coverages. Deliberate micro-mosaics had a longer strategy lifespan (17819 simulations) more often than rotations (15130 simulations). In comparisons where the simulations were the same length, micro-mosaics were generally more likely to maintain lower IR levels (5284 simulations) than rotations (1767) (Figure 7 top row). The sensitivity analysis (Figure 8), highlights that the benefit of deliberate micro-mosaics over rotations when considering improved biological and operational realism increased when female fitness costs increase. Figure 8 also highlights that deliberate micro-mosaics perform best when there is equal coverage of both insecticides. The benefit of deliberate micro-mosaics over rotations decreases when male fitness costs increase, cross resistance increases, dispersal increases or intervention coverage decreases.

**Figure 7.**
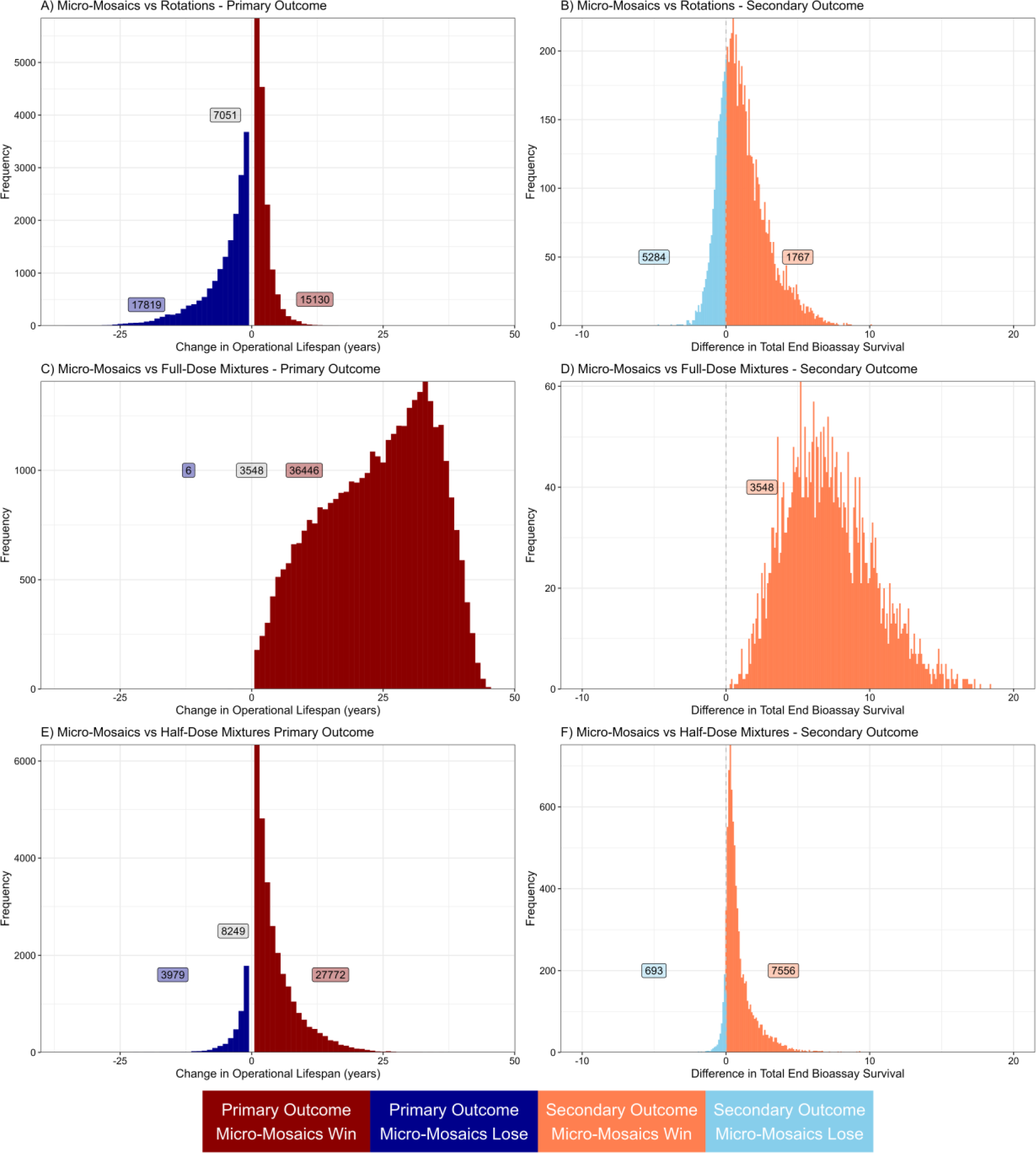
Scenario 1C Comparing Mismatched Coverage Micro-Mosaics versus Rotations (top), Full-Dose Mixtures (middle) and Half-Dose Mixtures (bottom) giving insecticides unique properties. Plots on the left evaluate on the primary outcome of the difference in strategy lifespan. Values to the left (dark blue) indicate the micro-mosaics strategy had a longer strategy lifespan, and values to the right (dark red) indicate the comparator strategy had a longer strategy lifespan. The numbers in each plot panel indicate the total of wins, losses and draws, with the grey boxes the number of draws in the strategy lifespan. Draws are evaluated on the secondary outcome of the peak bioassay survival shown in the plots on the right. Values to the left (light blue) indicate the micro-mosaics strategy had a lower peak bioassay survival, and values to the right (light red) indicate the comparator strategy had a lower peak bioassay survival.

**Figure 8:**
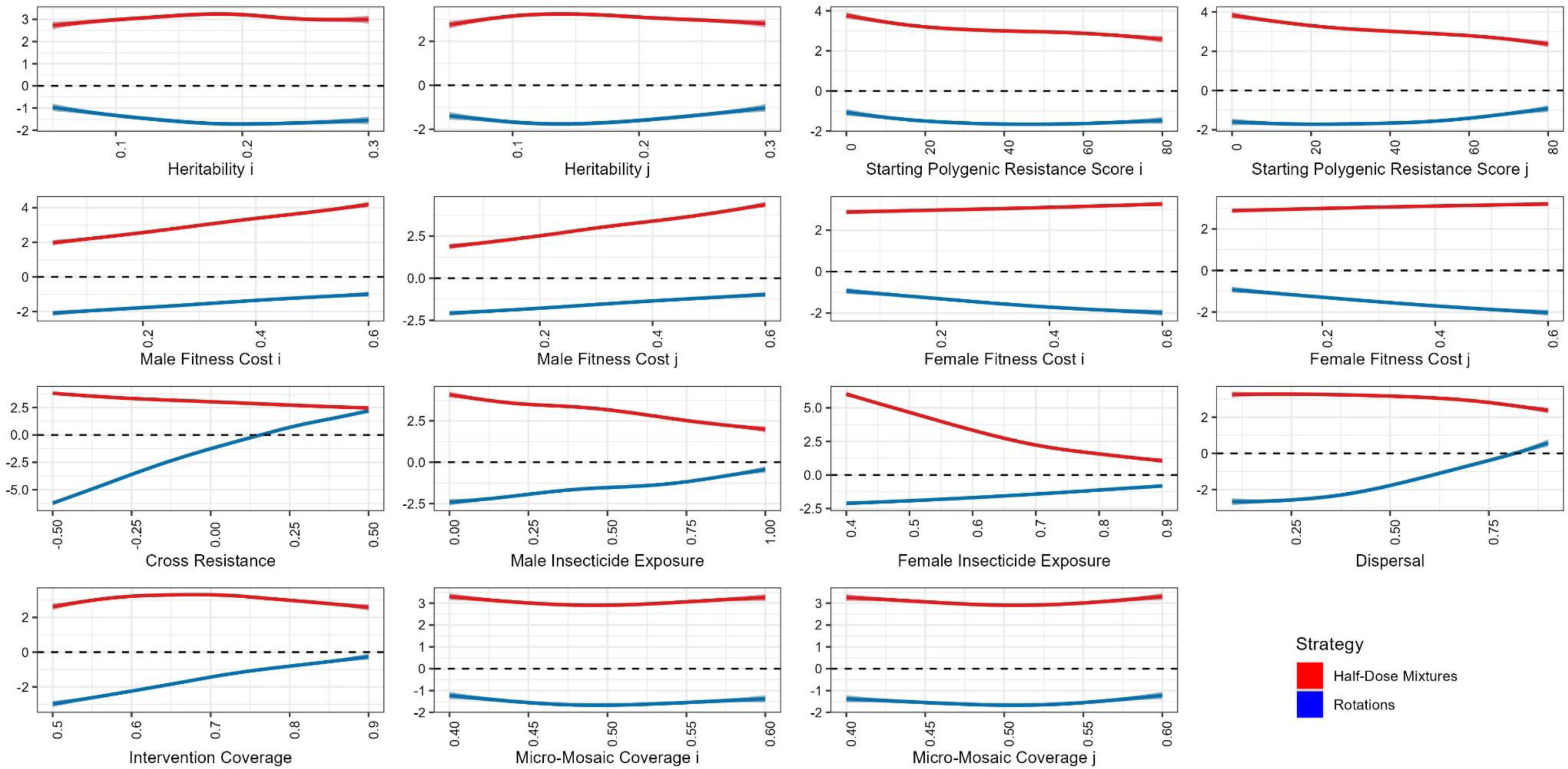
Sensitivity Analysis of Scenario 1C using Generalised Additive Models. Individual generalised additive models were fit for each randomly sampled parameter value used in Scenario 1C. The y axis is the difference in the strategy lifespan, negative numbers indicate the micro-mosaics performed better (below dashed line) and positive numbers indicate the comparator strategy performed better (above dashed line). The comparators are the half-dose mixtures (red) and rotations (blue). Full-dose mixtures were not included as they outperformed micro-mosaics regardless of the parameters used. The dashed line at 0 indicates the micro-mosaic and comparator strategy performed equally, and values above this line indicate the comparator strategy had a longer strategy lifespan, and values below the line indicate the micro-mosaic strategy had a longer strategy lifespan.

When comparing deliberate micro-mosaics against half-dose mixtures (Figure 7 middle row) it is seen that half-dose mixtures had a longer strategy lifespan (27772 simulations) more frequently than deliberate micro-mosaics (3979 simulations), and in comparisons where the strategy lifespans were equal, half-dose mixtures maintained lower levels of IR (7556 simulations) than for deliberate micro-mosaics (693 simulations).

When comparing deliberate micro-mosaics against full-dose mixtures (Figure 7 top bottom) full-dose mixtures had a longer strategy lifespan (36446 simulations) far more frequently than deliberate micro-mosaics (6 simulations), and in comparisons where the strategy lifespans were equal, full-dose mixtures maintained lower levels of IR (3548 simulations) than for deliberate micro-mosaics (0 simulations).

### 3.4. Scenario 1D: Evaluating Deliberate Micro-Mosaics with Insecticide Decay

Scenario 1D (Table 4) extended the simulations to include insecticide decay. The important question is whether the inclusion of insecticide decay into the simulations changes the qualitative outcome (which strategy performs best) and the quantitative advantage of the strategies.

When comparing deliberate micro-mosaics against rotations (Figure 9, top row), it is seen that when there is no cross resistance or negative cross resistance the micro-mosaic strategy generally performs best (regardless of the insecticide decay rates), and when there is positive cross resistance the rotation strategy performs best (again regardless of the insecticide decay rates). This is a qualitatively similar outcome to scenario 1A.

**Figure 9:**
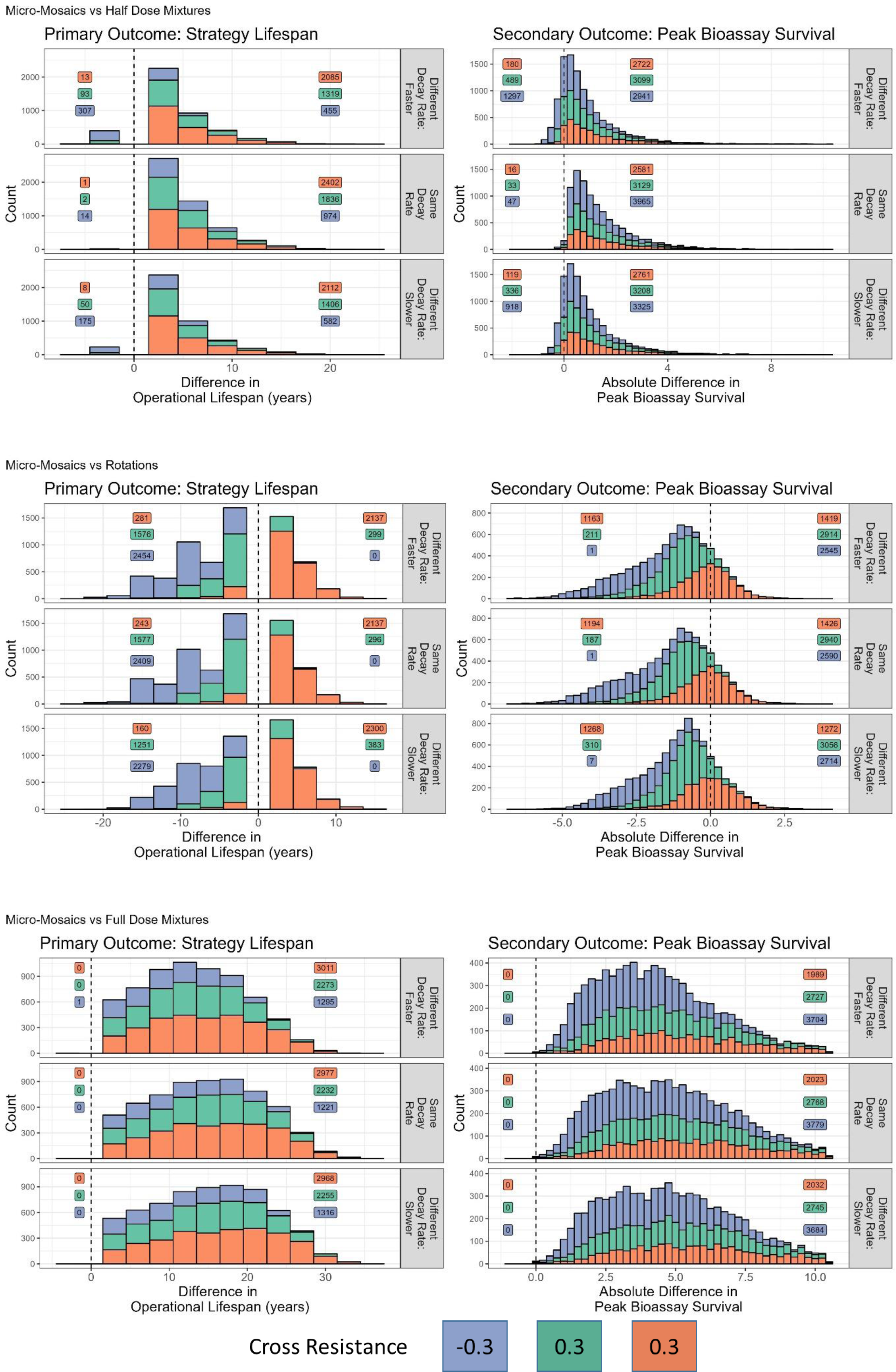
Scenario 1D Comparing Micro-Mosaics versus Rotations (top), Half-Dose Mixtures (middle) and Full-Dose Mixtures (bottom) when including insecticide decay. Plots on the left evaluate on the primary outcome of the difference in strategy lifespan, with values to the left indicating micro-mosaics had the longer strategy lifespan and values to the right indicate the comparator strategy had the longer strategy lifespan. Each plot is stratified by the base decay rates of the insecticides. For all panels insecticide *j* had a base decay rate of 0.015. Different decay rate faster (top panels) = insecticide *i* had a base decay rate of 0.025. Same decay rate (middle panels) = insecticide *i* had a base decay rate of 0.015. Different decay rate slower (bottom panels) = insecticide *i* had a base decay rate of 0.005. Draws are evaluated on the secondary outcome of the peak bioassay survival shown in the plots on the right. Values to the left indicate the micro-mosaics strategy had a lower peak bioassay survival, and values to the right indicate the comparator strategy had a lower peak bioassay survival. The colours of the bars indicate the degree of cross resistance between the insecticides, blue = negative (-0.3), red = positive (0.3) and green = no cross resistance (0). Numbers in the boxes in each panel indicate the number of wins/losses.

When comparing deliberate micro-mosaics against half-dose mixtures (Figure 9, middle row) it is seen that regardless of the level of cross resistance or rates of insecticide decay, the half-dose mixtures generally outperform the deliberate micro-mosaics. However, unlike in comparisons not including insecticide decay (Figure 3, middle row), there are now simulations where the deliberate micro-mosaic strategy outperforms half-dose mixtures, occurring mainly when there is negative cross resistance and the insecticides decay at different rates.

When comparing deliberate micro-mosaics against full-dose mixtures (Figure 9, bottom row) it is seen that regardless of the level of cross resistance or rates of insecticide decay, the full-dose mixtures outperform the deliberate micro-mosaics. This is a qualitatively identical result to comparisons not including insecticide decay (Scenario 1A, Figure 3, bottom row); however, the benefit of the mixture reduces, giving a 10-25 year longer strategy lifespan rather than 10-40 years seen when not including insecticide decay. Sensitivity analysis (Figure 10), highlights the comparative the parameter space where the micro-mosaics had an advantage over either rotations or half-dose mixtures.

**Figure 10:**
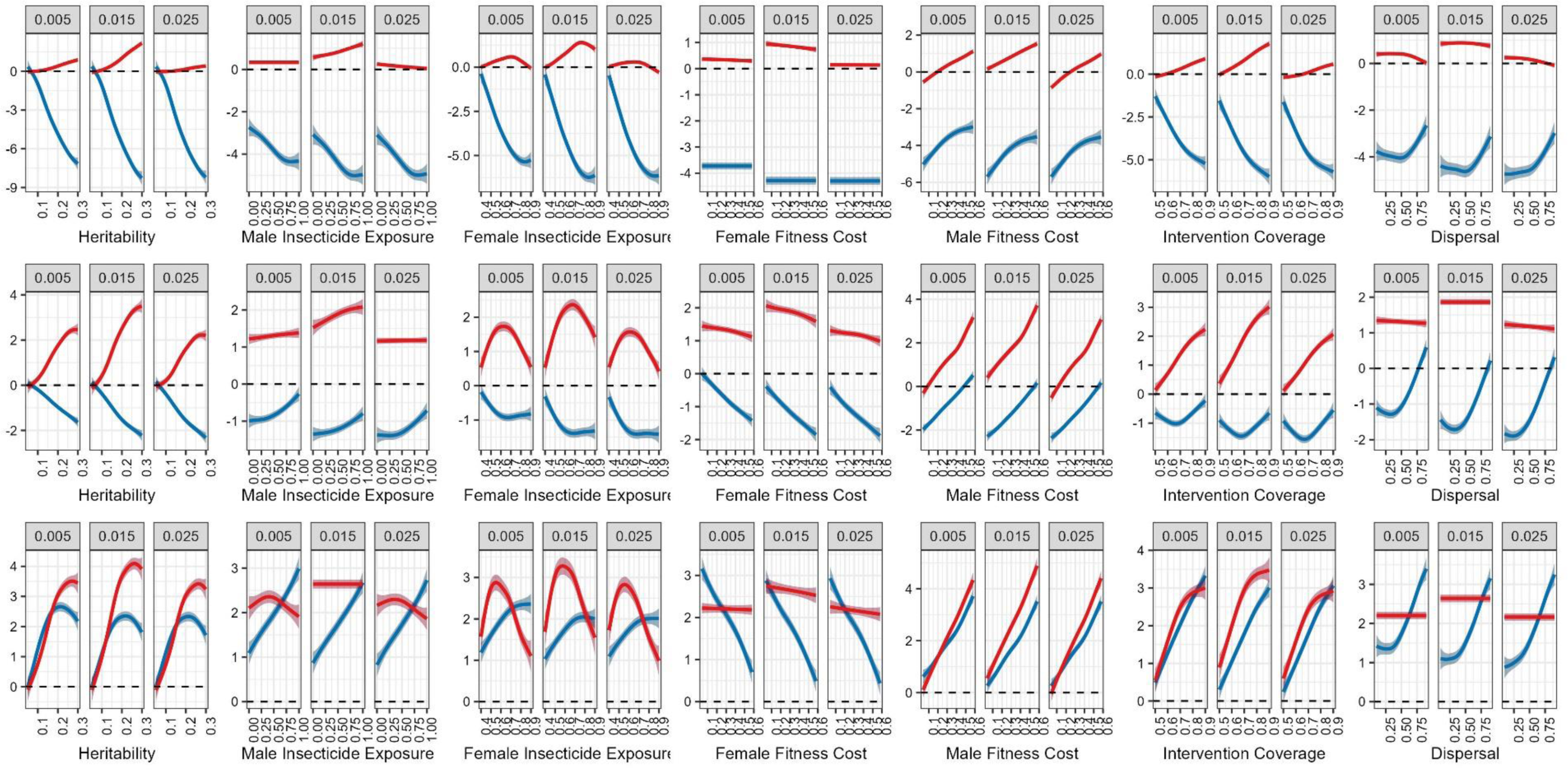
Generalised Additive Models for Scenario 1D. Top row is for negative cross resistance (α_*IJ*_ = -0.3), middle row is no cross resistance (α_*IJ*_ = 0.3), and bottom row is for positive cross resistance (α_*IJ*_ = 0.3). The y axis is the difference in Strategy Lifespan (in years) between the micro-mosaics strategy and the corresponding comparator strategies of rotations (blue) and half-dose mixtures (red). The full-dose mixture strategy was not included due to outperforming the micro-mosaics simulations for all simulations. Each plot is stratified by the base decay rate of insecticide *i*. The dashed line at 0 indicates the micro-mosaic and comparator strategy performed equally, and values above this line indicate the comparator strategy had a longer strategy lifespan, and values below the line indicate the micro-mosaic strategy had a longer strategy lifespan.

### 3.5. Scenario 2: Accidental Micro-Mosaics of Standard and Mixture ITNs

Scenario 2 (Table 5) concerned the deployment of mixture ITNs (either full-dose (Figure 11) or half-dose mixtures (Figure 12)) with standard-ITNs as accidental micro-mosaics. This was compared against both only mixture ITNs being deployed (expected upper benchmark) and only standard (pyrethroid-only) ITNs being deployed (expected lower benchmark) to understand the IRM implication of this occurring.

**Figure 11:**
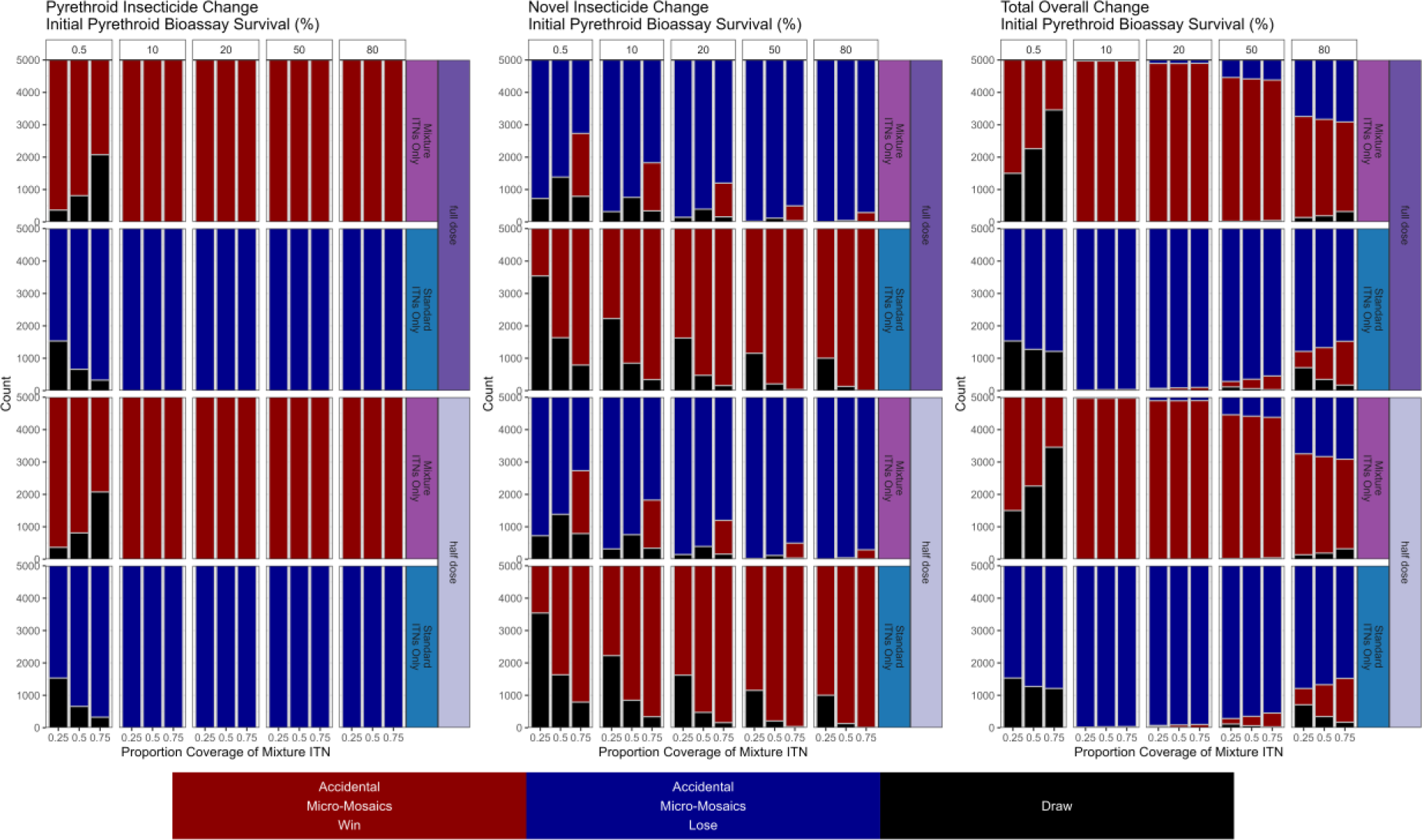
Scenario 2 - Implication of Accidental Micro-Mosaics. These mixture ITNs are deployed in a micro-mosaic with pyrethroid-only ITNs (insecticide *i*) which are at full-dose. These mixtures are compared over their proportional coverage of the mixtures (panels top-bottom) and the implication of pre-existing resistance to the pyrethroid (panels left-right). Left plots compares the change in bioassay survival for the pyrethroid insecticide. Middle plots compare the change in bioassay survival for the novel insecticide. Right plots compare the total bioassay survival change which is the change in bioassay survival for the pyrethroid insecticide plus the change in bioassay survival for the novel insecticide.

**Figure 12:**
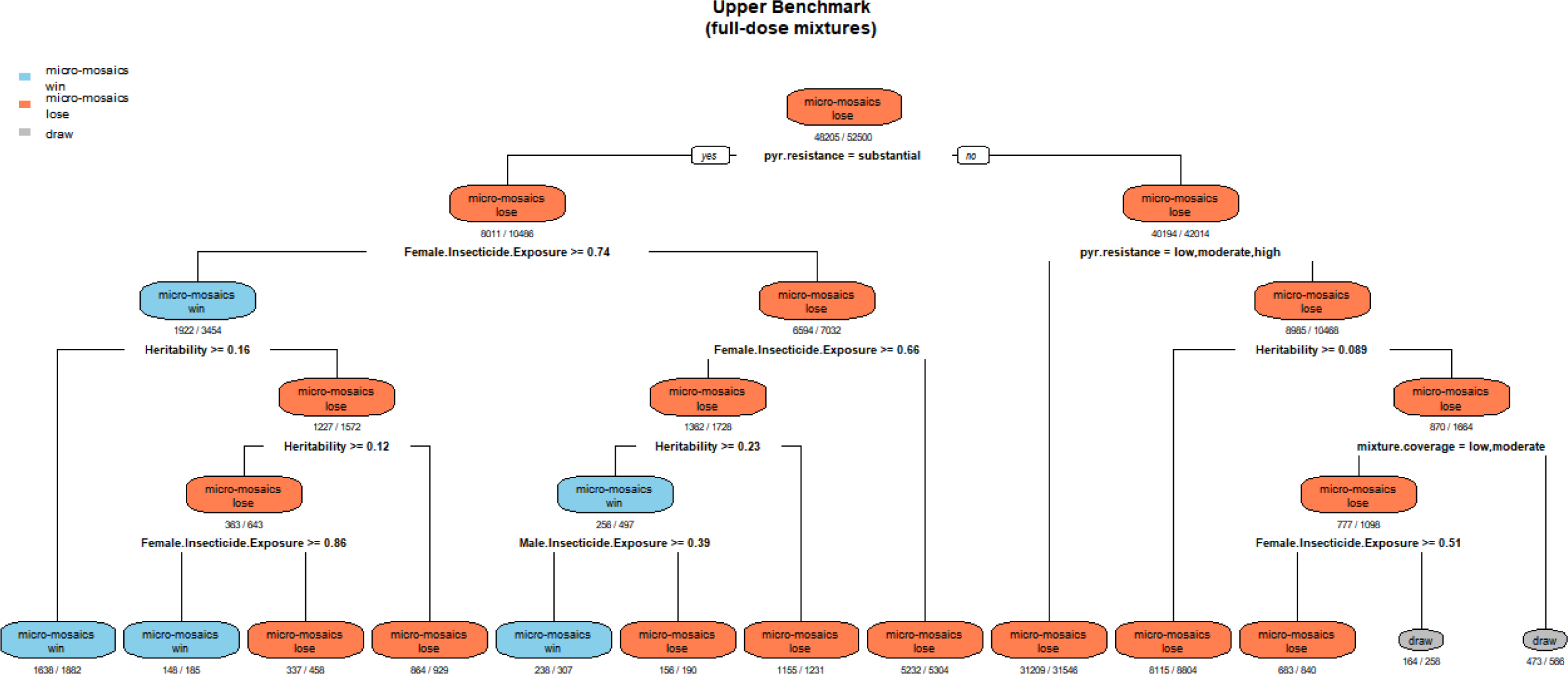
Regression Classification Tree Comparing Accidental Micro-Mosaics (with full-dose mixtures) versus Full-Dose Mixture ITNs Only. The outcome was the difference in the total change in resistance measured as bioassay survival, whereby micro-mosaics could end up with less total change (winning), more total change (losing) or the same total change (drawing). Pyr.resistance is the initial amount of resistance to the pyrethroid insecticide measured as bioassay survival and is classified as none (0.5%), low (10%), moderate (20%), high (50%) and substantial (80%). The term mixture.coverage is the proportion of the coverage which is mixture (*c*_*ij*_), with low = 0.25, moderate = 0.75 and high = 0.75. The colours indicate which strategy performed best, with blue = micro-mosaics win, red = micro-mosaics lose and grey = draw. The numbers underneath each leaf of the tree indicate the number of observations (dominator) and the number of observations with the corresponding outcome (numerator). The predictive accuracy of the model was 95.78%.

When considering full-dose mixtures used in an accidental micro-mosaic versus only full-dose mixtures, this leads to greater IR to the pyrethroid insecticide (Figure 11, top-left) but also lower IR to the novel insecticide (Figure 11, top-middle). The result is generally there being a lower amount of IR in total for the mixture only deployments (Figure 11, top-right). This indicates the presence of pyrethroid-only ITNs reduces the benefit of using full-dose mixtures. However, when there is already substantial IR to the pyrethroid there are occasions when the accidental micro-mosaic performs better overall. Sensitivity analysis highlights accidental micro-mosaics outperform mixtures only (from an IRM perspective) when there is substantial resistance to the pyrethroid (80% bioassay survival), the resistance traits have high heritabilities, and the female insecticide exposure is high (Figure 13).

**Figure 13:**
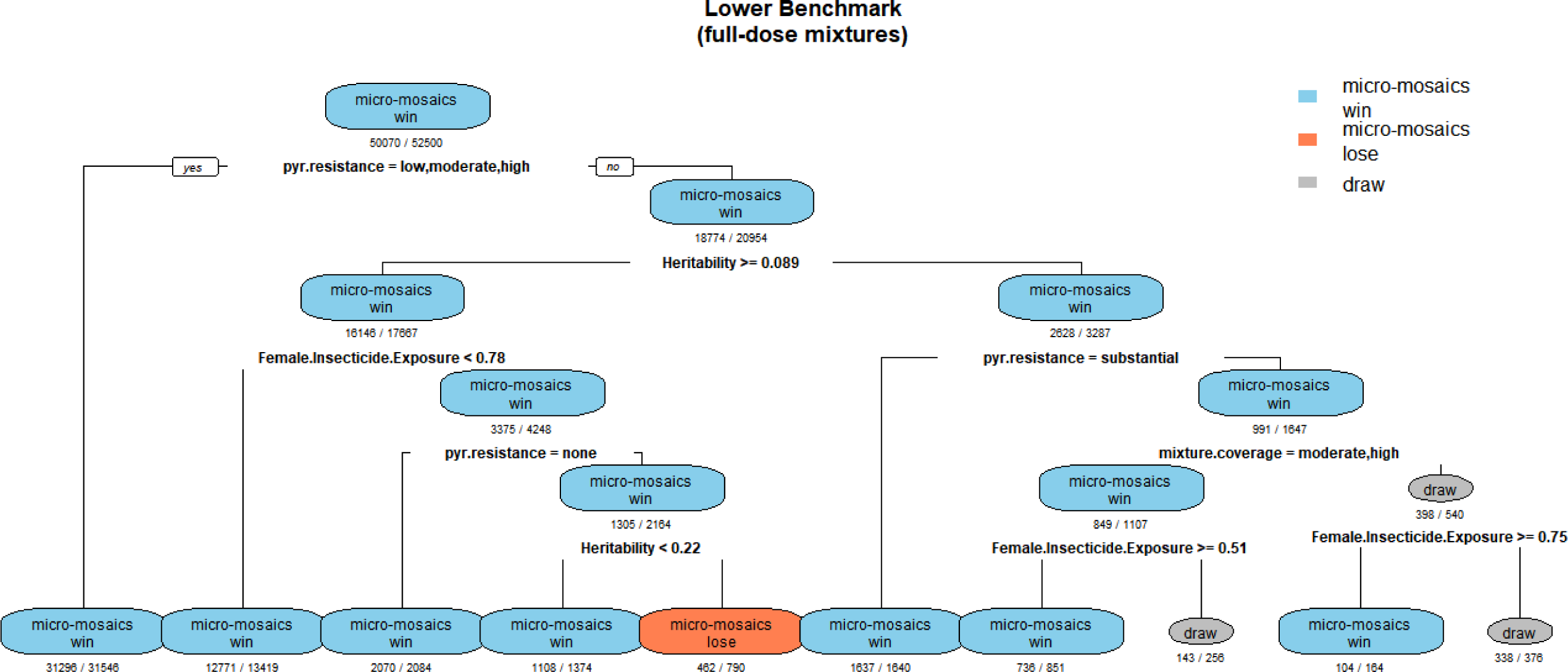
Regression Classification Tree Comparing Accidental Micro-Mosaics (with full-dose mixtures) versus Standard ITNs Only. The outcome was the difference in the total change in resistance measured as bioassay survival, whereby micro-mosaics could end up with less total change (winning), more total change (losing) or the same total change (drawing). Pyr.resistance is the initial amount of resistance to the pyrethroid insecticide measured as bioassay survival and is classified as none (0.5%), low (10%), moderate (20%), high (50%) and substantial (80%). The term mixture.coverage is the proportion of the coverage which is mixture (*c*_*ij*_), with low = 0.25, moderate = 0.75 and high = 0.75. The colours indicate which strategy performed best, with blue = micro-mosaics win, red = micro-mosaics lose and grey = draw. The numbers underneath each leaf of the tree indicate the number of observations (dominator) and the number of observations with the corresponding outcome (numerator). The predictive accuracy of the model was 96.62%.

When the full-dose mixture micro-mosaics are compared against pyrethroid ITNs only, it is seen that there is a decrease in the IR to the pyrethroid for the accidental micro-mosaics. There is an increase in the level of IR to the novel insecticide when compared to the standard ITN only, which is expected as the novel insecticide is not deployed. When looking at the overall change in the total IR, the accidental micro-mosaics outperform the pyrethroid-only ITNs except at very high levels of initial IR to the pyrethroid (Figure 11). Sensitivity analysis highlights the pyrethroid only ITNs outperform the accidental micro-mosaics if there is already substantial resistance to the pyrethroids (80% bioassay survival), female insecticide exposure is very high, and the heritability of the resistance traits is very high (Figure 13).

When considering half-dose mixtures used in an accidental micro-mosaic versus only half-dose mixtures, it can be seen this leads to greater IR to the pyrethroid insecticide but also lower IR to the novel insecticide. The result is there generally being a lower amount of IR in total for the mixture only deployments. This indicates the presence of pyrethroid-only ITNs reduces the benefit of using full-dose mixtures. However, when there is already substantial IR to the pyrethroid there are occasions when the micro-mosaic performs better overall occasionally substantially (Figure 11). Sensitivity analysis highlights the accidental micro-mosaics outperform the mixtures only when the resistance to the pyrethroid is high (greater than 50% bioassay survival), the heritability of the traits is very high, and the female insecticide exposure is very high (Figure 14).

**Figure 14:**
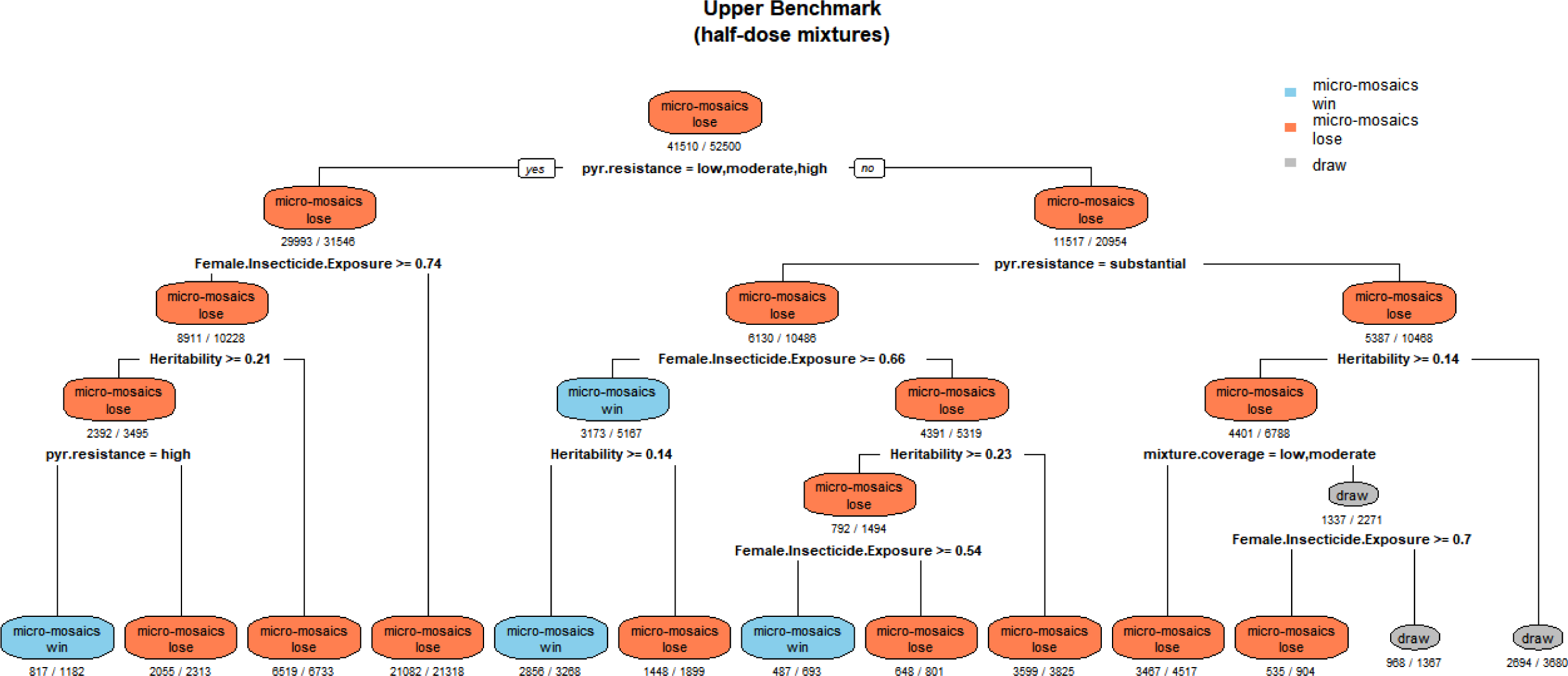
Regression Classification Tree Comparing Accidental Micro-Mosaics (with half-dose mixtures) versus Half-Dose Mixture ITNs Only. The outcome was the difference in the total change in resistance measured as bioassay survival, whereby micro-mosaics could end up with less total change (winning), more total change (losing) or the same total change (drawing). Pyr.resistance is the initial amount of resistance to the pyrethroid insecticide measured as bioassay survival and is classified as none (0.5%), low (10%), moderate (20%), high (50%) and substantial (80%). The term mixture.coverage is the proportion of the coverage which is mixture (*c*_*ij*_), with low = 0.25, moderate = 0.75 and high = 0.75. The colours indicate which strategy performed best, with blue = micro-mosaics win, red = micro-mosaics lose and grey = draw. The numbers underneath each leaf of the tree indicate the number of observations (dominator) and the number of observations with the corresponding outcome (numerator). The predictive accuracy of the model was 89.8%.

When the half-dose mixture accidental micro-mosaics are compared against pyrethroid ITNs only, it is seen that there is a decrease in the IR to the pyrethroid for the accidental micro-mosaics. There is an increase in the level of IR to the novel insecticide when compared to the standard ITN only, which is expected as the novel insecticide is not deployed. When looking at the overall change in the total IR, the accidental micro-mosaics outperform the pyrethroid-only ITNs except at very high levels of initial IR to the pyrethroid (Figure 11). Sensitivity analysis highlights the pyrethroid only ITNs outperform the accidental micro-mosaics if there is already substantial resistance to the pyrethroids (80% bioassay survival), female insecticide exposure is high, and the heritability of the resistance traits is high (Figure 15).

**Figure 15:**
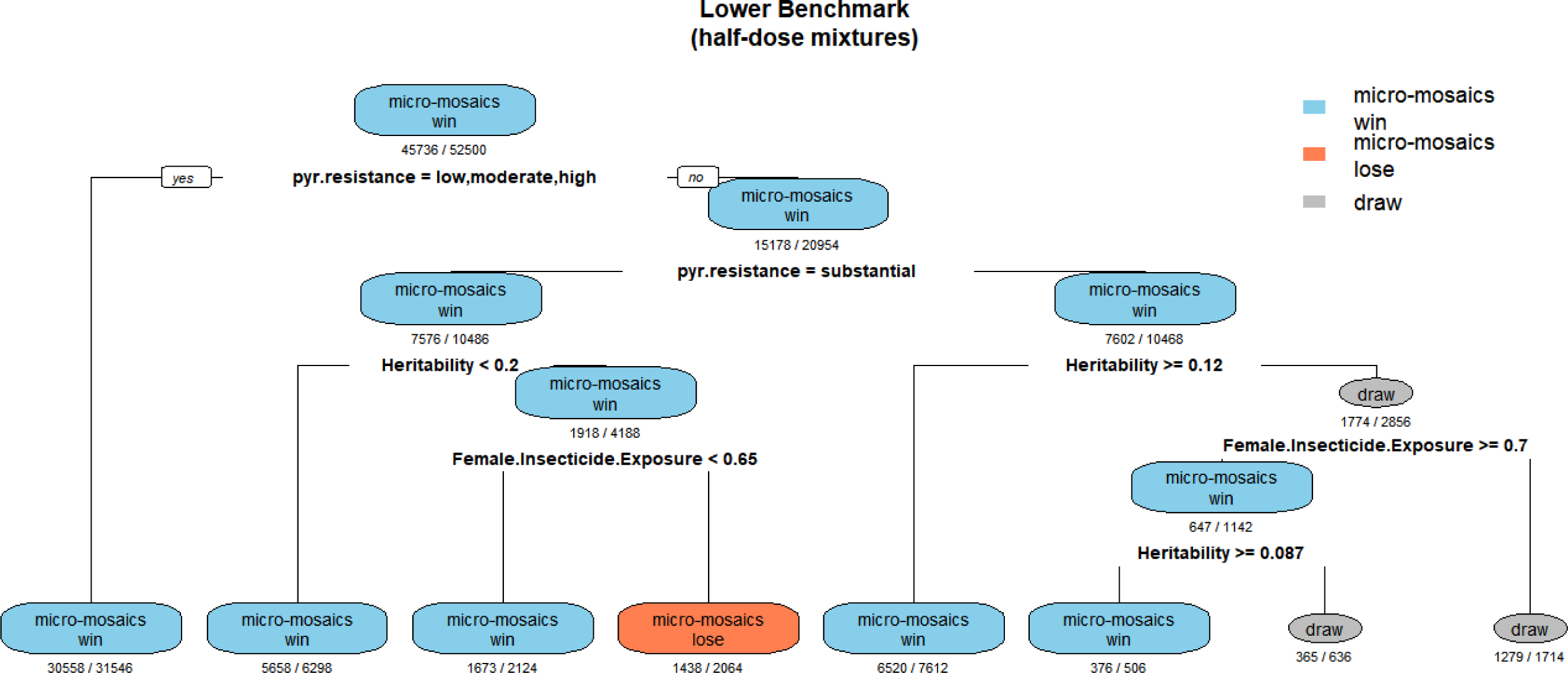
Regression Classification Tree Comparing Accidental Micro-Mosaics (with half-dose mixtures) versus Standard ITNs Only. The outcome was the difference in the total change in resistance measured as bioassay survival, whereby micro-mosaics could end up with less total change (winning), more total change (losing) or the same total change (drawing). Pyr.resistance is the initial amount of resistance to the pyrethroid insecticide measured as bioassay survival and is classified as none (0.5%), low (10%), moderate (20%), high (50%) and substantial (80%). The colours indicate which strategy performed best, with blue = micro-mosaics win, red = micro-mosaics lose and grey = draw. The numbers underneath each leaf of the tree indicate the number of observations (dominator) and the number of observations with the corresponding outcome (numerator). The predictive accuracy of the model was 91.36%.

## 4. Discussion

As more insecticidal products become available, there is an increase in the number and complexity of IRM strategies to consider. One such strategy is the “micro-mosaic” strategy which involves distributing different insecticides to different households (Levick et al., 2017) in the same village with the hope of gaining a “temporal mixture” (Jones et al., 2023). This is clearly a logistically complex IRM strategy to implement and therefore demonstrating a theoretical benefit of micro-mosaics should be required before considering field trials which are expensive and resource intensive (Tabashnik, 1986). A thorough evaluation of the micro-mosaic strategy has been presented, including the implication of cross resistance, higher resistance settings, insecticide decay and the implication of accidental micro-mosaics for mixture ITN deployments with standard (pyrethroid-only) ITNs.

### 4.1. Evaluation of Micro-Mosaics as a Deliberate IRM strategy

Comparisons between deliberate micro-mosaics, rotations, and half-dose mixtures (such that the “amount of insecticide” deployed at any timepoint is consistent between all strategies), show there is generally very limited benefit in choosing one strategy over another, with strategies generally adding an extra 5 years to the strategy lifespan of the insecticides. (Figures 3, 5, 7, 9) regardless of the inclusion of higher resistance, insecticide decay or cross resistance. Full-dose mixtures perform well due to have a greater total amount of insecticide deployed at any time. The general poor performance of the micro-mosaics strategy compared to full-dose mixtures is consistent with an evaluation using assuming a monogenic basis of resistance (Jones et al., 2023). Different models with different assumptions reaching similar conclusions improves confidence in theoretical investigations evaluating the performance of IRM strategies.

It could be argued that, in practice, the best comparator for deliberate micro-mosaics is the rotations strategy (top panels of Figures 3, 5, 7, 9). This allows for the two insecticides to be manufactured by different companies and are deployed as separate formulations. The use of mixtures necessitates both insecticides to be manufactured by the same company, on the provision they can form a chemically stable mixture formulation. The putative theoretical advantage of deliberate micro-mosaics was to gain a “temporal mixture”, potentially of a form which would not be physically made, either for chemical or commercial reasons.

### 4.2. Implication of Accidental Micro-Mosaics of Standard and Mixture ITNs

Our results indicate the presence of standard (pyrethroid-only) ITNs in an accidental micro-mosaic with mixture ITNs generally reduces the IRM potential of the mixture ITNs (Figure 11 and 12) compared to if mixture ITNs were deployed alone. Perhaps an important question to answer would be how frequently do micro-mosaics currently accidentally occur in the field? While currently this would likely be an accidental micro-mosaic of standard pyrethroid nets of which cross resistance is likely to be substantial (Moyes et al., 2021), this has been found to be occurring in cluster-RCTs evaluating next-generation mixture nets (Accrombessi et al., 2023; Mosha et al., 2022) indicating this will likely be an issue come wider rollout of mixture ITNs.

This issue of accidental micro-mosaics is further complicated by the issue of the simultaneous use of mixture nets containing pyrethroid+chlorfenapyr and synergist nets containing pyrethroid+piperonyl butoxide (PBO). This is because chlorfenapyr is a pro-insecticide that must be converted into the active form my P450 enzymes (Black et al., 1994). Unfortunately, these P450 enzymes are inhibited by PBO which may well reduce the activity of chlorfenapyr. PBO however increases effectiveness of pyrethroid by inhibiting the P450s which cause metabolic resistance to pyrethroid (David et al., 2013). The implication of this was recently tested in experimental hut trials (Syme et al., 2023), where it was found that the presence of both chlorfenapyr and PBO nets in the same hut (and therefore encountered in the same gonotrophic cycle) reduced their overall kiIling efficacy. We would extend these concerns regarding reduced killing to being less effective for IRM also.

### 4.3. Caveats of Micro-Mosaic Evaluation

While a technically challenging strategy to implement, if technical/logistical hurdles around the deployment of deliberate micro-mosaics can be overcome, then this strategy does offer benefits which our simulations did not directly account for. These caveats may either improve, or reduce, the IRM potential of micro-mosaics,.

#### 4.3.1. Caveat: Time to Market

When designing simulations to evaluation IRM strategies, we attempt to make as best direct comparisons between the strategies. That is having a fixed number of insecticides involved in each strategy. For example, our micro-mosaic consisted of two insecticides, and the corresponding rotation simulations also consisted of two insecticides. However, there is no reason to not also allow for rotations to occur within a micro-mosaic too. Such that in the timeframe for deploying two insecticides in a standard rotation, four different insecticides would have been deployed in a rotating deliberate micro-mosaic (insecticide A and B in deployment round 1, insecticide C and D in deployment round 2). This achieves two things, first halving the selection pressure again, but also increasing the frequency of deployments such that products can reach the field sooner. For example, in a four-ITN rotation, the fourth ITN would not be deployed until 9 years had passed, whereas with micro-mosaics this would be only after 3 years. A general assumption is that an increased number of insecticides and diversity of insecticides is beneficial for IRM. The model only currently allows for a maximum of two insecticides to be deployed at any time point and if logistical hurdles involved deploying multiple separate insecticides can be overcome, there is no reason not to extend to include a greater number of insecticides in the micro-mosaic.

#### 4.3.2. Caveat: Implementation Complexity

Deliberate micro-mosaics would clearly be a logistically challenging IRM strategy to implement, requiring multiple supply chains. This increases the opportunity for a procurement/supply/logistics failure, which could mean that only half the required amount of insecticide is available for use. Clearly, with more complex insecticide deployments there are greater opportunities for operational issues to occur. Levick et al (2017) included a “correct deployment parameter” in their model to allow for failures in correct deployments for mixtures (e.g., only one mixture partner is deployed). For deliberate micro-mosaics this could occur in one of two ways. First, is the failure to procure and/or deploy one of the insecticides (e.g., insecticide *i*) used in the deliberate micro-mosaic, without replacing the missing coverage with additional of the other insecticide (e.g., insecticide *j*). By reducing the overall selection pressure this would make the strategy appear better from an IRM stand-point, however this would be at the detriment of providing less control. Second, is the failure to procure and/or deploy one of the insecticides (e.g., insecticide *i*) used in the micro-mosaic, but with the missing coverage replaced by insecticide *j* and this would turn the strategy more into the rotation and/or sequence strategy with monotherapy deployments.

## Conclusion

Unfortunately, the deliberate use of micro-mosaics was found to not be beneficial IRM strategy, especially when compared against its most obvious comparator of rotations. Micro-mosaics occurring unintentionally due to multiple distribution channels may reduce the overall benefit of mixture nets.

## Author Contributions

- NPH: Data Curation, Software, Formal Analysis, Investigation, Methodology, Writing – Original Draft Preparation
- IMH: Investigation, Methodology, Writing – Review & Editing, Supervision

## Data Availability Statement

Model code for the running and analysis of simulations is available from the github respository: https://github.com/NeilHobbs/polysmooth and the corresponding author upon request.

## Acknowledgements

We would like to acknowledge the advice of David Weetman for help grounding the model in biological and operational relevance. We would also like to acknowledge the Vector Informatics and Genomics group at the Liverpool School of Tropical Medicine for their general critiques of the model methodology.

